# RNAGenesis: A Generalist Foundation Model for Functional RNA Therapeutics

**DOI:** 10.1101/2024.12.30.630826

**Authors:** Zaixi Zhang, Ruofan Jin, Linlin Chao, Guangxue Xu, Yikun Zhang, Guowei Zhou, Di Yin, Yingqing Guo, Yaqi Fu, Yukang Yang, Kaixuan Huang, Xiaotong Wang, Junze Zhang, Yujie Yang, Qirong Yang, Ziyao Xu, E Weinan, Ruhong Zhou, Xiaoming Zhang, Mengdi Wang, Le Cong

## Abstract

RNA molecules are central to gene regulation, catalysis, and molecular recognition, and offer broad opportunities for therapeutic applications. However, uncovering their complex sequence, structure, and function relationships, particularly for non-coding RNAs, remains a formidable challenge. Here, we introduce *RNAGenesis*, a Generalist RNA foundation model that integrates sequence representation, structural prediction, and de novo functional design within a single generative framework. Trained on diverse clustered non-coding RNAs, RNAGenesis leverages a BERT-style encoder, query-based latent compression, and a diffusion-guided decoder enhanced by inference-time alignment with gradient guidance and beam search strategies. Through comprehensive evaluations, RNAGenesis achieves state-of-the-art performance on 11 of 13 tasks in the BEACON benchmark and surpasses structure-aware models in inverse folding, 3D structure prediction, and de novo structure design. We further introduce *RNATx-Bench*, a dedicated benchmark for RNA therapeutics comprising over 100,000 experimentally validated sequences. RNAGenesis demonstrates strong predictive performance across ASOs, siRNAs, shRNAs, circRNAs, and untranslated region (UTR) variants. Furthermore, RNAGenesis enables functional RNA design, including aptamers targeting IGFBP3 and structurally constrained sgRNA scaffolds. Wet-lab validation confirms aptamer binding with *K*_*D*_ values as low as 4.02 nM and up to 2.5-fold improvement in editing efficiency across CRISPR-Cas9, base editing, and prime editing systems. These results position RNAGenesis as a next-generation general-purpose RNA foundation model with broad utility for computational modeling and experimental therapeutic design.

## Main

RNA molecules play fundamental roles across a wide spectrum of biological processes, extending far beyond their canonical function as messengers in the central dogma of molecular biology. In addition to ribosomal RNA (rRNA) and transfer RNA (tRNA), which coordinate protein synthesis, a rapidly expanding repertoire of non-coding RNAs (ncRNAs) orchestrates diverse regulatory and catalytic functions. For example, microRNAs (miRNAs) and long non-coding RNAs (lncRNAs) are central to complex gene regulatory networks^1, 2^, small interfering RNAs (siRNAs) mediate post-transcriptional gene silencing^3^, and CRISPR-associated RNAs, such as single guide RNAs (sgRNAs), enable precise genome editing^4^. These functional roles are governed by the intricate interplay between RNA sequence, higher-order structure, and biological activity, which in turn shapes cellular behavior and contributes to the etiology of many diseases. Deciphering these sequence–structure–function relationships is thus critical for advancing our understanding of RNA biology, accelerating RNA-based therapeutics, and enabling rational RNA design in synthetic biology.

In response to this need, a growing number of RNA foundation models have emerged, leveraging large-scale RNA sequence datasets to learn evolutionarily meaningful representations for diverse downstream tasks^5–11^. Most of these models adopt encoder-only Transformer architectures trained with masked language modeling (MLM) objectives. Notable examples include RNA-FM^6^ and RiNALMo^10^, both of which are BERT-style models trained on the RNACentral database^12^ and demonstrate strong generalization to unseen RNA families. RNAErnie^13^ incorporates RNA motifs as biological priors and introduces RNA-type tokenization (e.g., miRNA, lncRNA) to guide fine-tuning. AIDO.RNA^11^ scales these advances to larger model sizes (1.6B parameters), achieving new benchmarks across a wide range of RNA tasks. Several specialized models have also been proposed: UTR-LM^7^ focuses on untranslated regions (UTRs), revealing regulatory motifs that govern gene expression, while CodonBERT^14^ targets codon-level modeling to capture codon usage bias and its impact on translation efficiency.

While recent RNA foundation models have shown promise in representation learning, they are largely limited in generative and structural capabilities, restricting their utility in programmable RNA design and therapeutic development. Here, we introduce **RNAGenesis**, a unified foundation model with 1 billion parameters that integrates RNA sequence understanding, *de novo* design, and 3D structure prediction within a single framework. Despite its compact size, RNAGenesis combines a pretrained encoder, a latent diffusion decoder, and multi-modal fusion modules to deliver strong performance across a broad range of RNA modeling and design tasks. On 13 RNA tasks from the BEACON benchmark^15^, RNAGenesis ranks first on 11, demonstrating broad and robust capabilities. To systematically evaluate therapeutic applications, we further introduce **RNATx-Bench**, a dedicated benchmark comprising over 100,000 experimentally validated RNAs across six major modalities: antisense oligonucleotides (ASOs), siRNAs, shRNAs, circRNAs, aptamers, and human UTR variants. RNAGenesis outperforms leading foundation models such as RNA-FM^6^ and Evo2^16^ across predictive and generative tasks, including highest correlation scores for ASO efficacy on clinically relevant genes, AUROC > 0.8 in siRNA prediction, and over 10% correlation gains on shRNA benchmarks. It also generates aptamers with improved GC content, structural similarity, and free energy compared to SELEX-derived sequences. For UTR variants, RNAGenesis accurately classifies pathogenicity and regulatory effects, ranking among the top models on ClinVar, IRD, and MPRA datasets. Beyond sequence, RNAGenesis incorporates pretrained structural modules to support structure-conditioned tasks such as inverse folding, 3D structure prediction, and de novo structure generation. It surpasses RhoDesign^17^ in inverse folding (57.1% recovery, 16.9 °A RMSD, TM-score 0.442), ranks second-best in structure prediction while being the fastest to infer, and produces more native-like folds across variable lengths (40–150 nt) with higher scTM scores and accurate local geometries. Together, these results establish RNAGenesis as a versatile and efficient platform for unified RNA modeling and programmable therapeutic engineering.

To advance programmable RNA design, we apply RNAGenesis to both engineer RNA aptamers for target protein binding and optimize sgRNA scaffolds for genome editing. By integrating latent diffusion with inference-time alignment—through gradient-based guidance and beam search—RNAGenesis generates sequences tailored for both structural stability and functional efficacy (Figure 4a–c). In wet-lab experiments, the designed **aptamers** achieved binding affinities as low as 4.02nM against the IGFBP3 target. For genome editing, 6 out of 24 designed sgRNA scaffolds outperformed the wild-type in a GFP reporter assay. At endogenous loci such as AAVS1 and B2M, RNAGenesis designs led to up to a 2-fold increase in knockout efficiency across varying dosage levels (Figure 4d). We next assessed whether these scaffolds generalize to more advanced and clinically relevant genome editing systems, including **base editing** and **prime editing**—technologies that enable precise single-nucleotide changes or small insertions/deletions without inducing double-strand breaks, offering transformative potential for correcting pathogenic mutations^18, 19^. Without any model retraining, RNAGenesis-designed scaffolds significantly improved editing outcomes in both systems. In cytosine base editing (CBE), the RGen-6 scaffold enhanced editing efficiency by over 2.5-fold. In prime editing—requiring more intricate RNA design—RGen-6 increased efficiency by up to 1.2-fold compared to the wild-type pegRNA (Figure 4h–l). These findings highlight the strong zero-shot generalization of RNAGenesis to diverse RNA-guided genome editing modalities. To elucidate the molecular basis of these improvements, we conducted structural analyses of RNAGenesis-designed scaffolds. Predictions using AlphaFold3^20^ and Boltz-2^21^ revealed that optimized scaffolds maintain the global secondary structure while exhibiting lower minimum free energy and enhanced thermodynamic stability. Notably, RGen-6 featured more stable G–C base pairing, a higher number of hydrogen bonds (56 versus 43), and improved MMPBSA binding free energy (–388 versus –336 kcal/mol) relative to the wild-type scaffold (Figure 4g,k,o). These structural enhancements likely underpin the improved editing efficiency observed across CRISPR, base editing, and prime editing platforms.

Together, these results establish RNAGenesis as a versatile and generalist RNA foundation model capable of designing functional RNA molecules across diverse therapeutic and genome engineering applications—including aptamer binding, CRISPR editing, and clinically relevant base and prime editing—with minimal adaptation.

### Unified Model for RNA Understanding and Design

In this work, we present **RNAGenesis**, a unified RNA foundation model that bridges RNA sequence understanding and *de novo* sequence design through latent diffusion^22^ (Figure 1a–c). RNAGenesis is pretrained on a large-scale corpus of RNA sequences from RNAcentral, covering a diverse range of RNA categories including mRNA, lncRNA, rRNA, tRNA, miRNA, snoRNA, and ribozymes. Designed for both efficiency and scalability, RNAGenesis is a compact foundation model with approximately 1 billion parameters. Its encoder follows a BERT-style Transformer architecture comprising 32 layers, augmented with hybrid N-gram tokenization and 1D convolutional layers to capture both single-nucleotide resolution and multi-scale contextual signals (Figure 1b). Further architectural details are provided in the Methods section.

**Figure 1.**
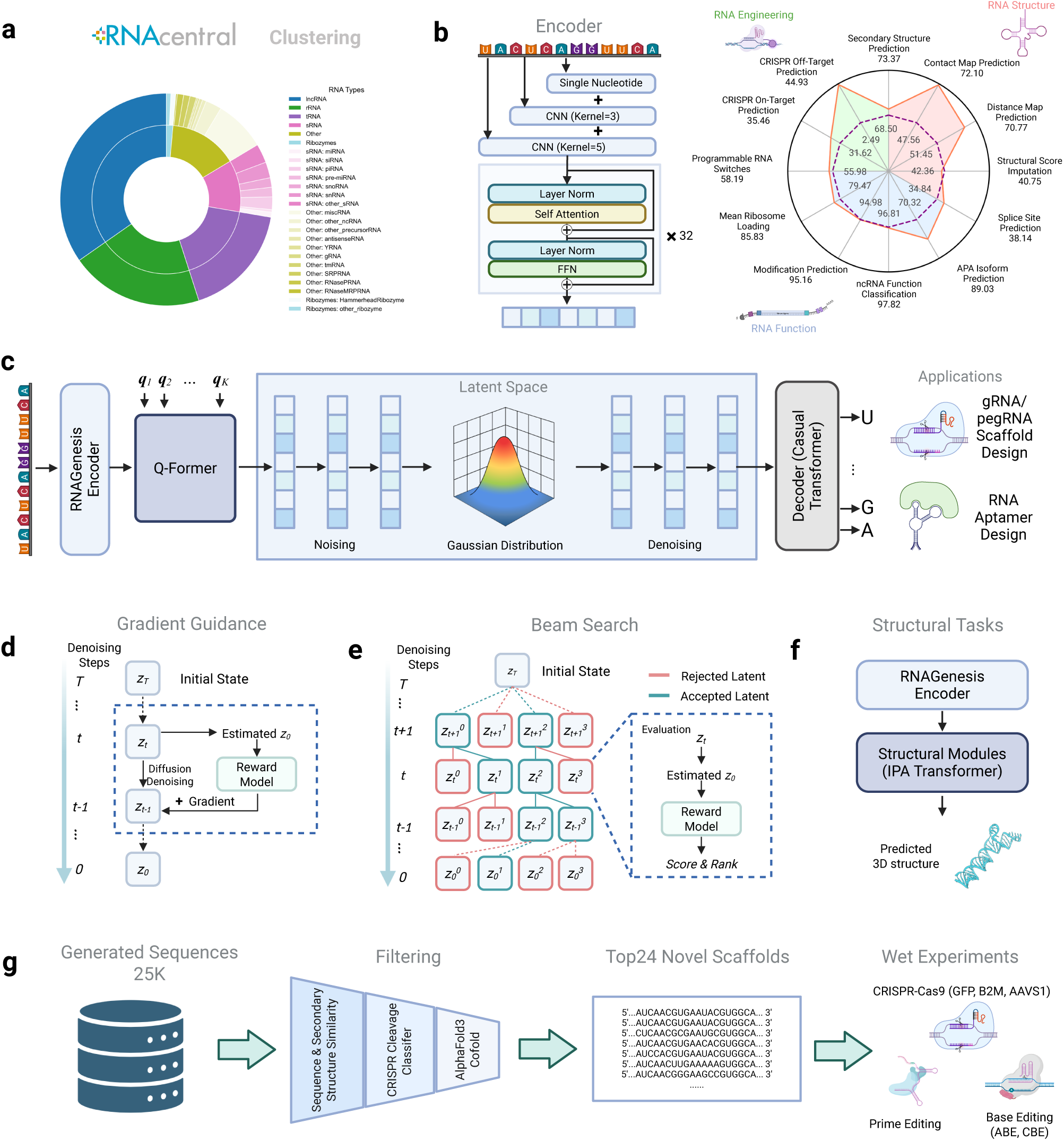
Overview of RNAGenesis: a Generalist RNA Foundation model unifies sequence understanding, sequence generation, and structural prediction. **(a)** RNAGenesis is trained on RNACentral with sequence clustering. The main RNA types include lnRNA, rRNA, tRNA, small RNA (sRNA), ribozymes, and other types. **(b)** The Encoder of RNAGenesis consists of Hybrid N-Gram Tokenization with single nucleotide representation and One-dimensional Convolutional Neural Networks (1D CNNs) and the 32-layer Transformer backbone. The input is an RNA sequence, and the resulting representation can be utilized for a range of downstream tasks, including structure, function, and engineering-related predictions. On the Beacon benchmark, RNAGenesis (orange) overperforms RNA-FM (purple) on nearly all tasks. **(c)** RNAGenesis uses a latent diffusion method for RNA sequence generation, such as *de novo* gene editing gRNA/pegRNA scaffold and RNA aptamer design. The output RNA representation from RNAGenesis Encoder further undergoes Q-Former transformation to obtain the latent representations. In the latent diffusion, RNAGenesis learns to denoise from the Gaussian Distribution, after which the final RNA sequence is generated using a Causal Transformer Decoder. **(d & e)**. Inference-time alignment with Gradient Guidance and Beam Search in the latent space. **(d)** Gradient Guidance deals with a differentiable reward model (e.g., trained neural networks for sgRNA cleavage classification). During each denoising step from *T* to 0, the reward model, the gradient is calculated and added to the current states based on the estimated state at *t* = 0 and the reward model;(e) Beam Search is used for a non-differentiable reward model (e.g., minimum free energy calculation). It works by seeking a better diffusion path over the denoising process, sampling *K* latents per beam, and possessing ***B*** beams for the next step. The Reward model scores and ranks the latents at *t*, accepts the top-ranked latents for further denoising steps, and rejects the others. (f) RNAGenesis can be equipped with structural modules (e.g., IPA Transformer) to tackle structural tasks such as 3D structure predictions. (g) Wet experiments on CRISPR-Cas9 (GFP, B2M, AAVS1), Prime Editing, and Base Editing are conducted to verify the effectiveness of RNAGenesis.

To compress high-dimensional sequence embeddings into a lower-dimensional latent space suitable for generation, RNAGenesis employs a Query Transformer (Q-former)^23^, which maps encoder outputs into fixed-length latent vectors. These vectors serve as inputs to a denoising diffusion model, trained to capture the distribution of functional RNA sequences in the latent space. The final RNA sequence is decoded autoregressively using a Causal Transformer decoder. This architecture enables powerful sequence generation capabilities, from *de novo* sgRNA and pegRNA scaffold design to aptamer generation and general RNA design.

To further improve generation quality, we introduce two inference-time optimization strategies: Gradient Guidance and Beam Search (Figure 1 d–e). Gradient Guidance involves a differentiable reward model (e.g., an sgRNA cleavage classifier) that provides gradients during each denoising step, nudging the latent vector toward desired properties. In contrast, Beam Search is used with non-differentiable objectives such as minimum free energy (MFE), selecting and refining multiple candidate paths in latent space to maximize sequence quality.

Beyond sequence modeling, RNAGenesis can be extended to structural tasks through multi-modal integration with pretrained structure-aware modules (Figure 1 f). For example, we pair the RNAGenesis encoder with the IPA Transformer from RNA-FrameFlow^24^ to enable downstream structural predictions, including RNA 3D structure prediction and *de novo* structure generation. This multi-modality makes RNAGenesis a comprehensive framework for both RNA understanding and design.

Finally, to validate the real-world applicability of RNAGenesis, we conduct wet-lab experiments (Figure 1 g), applying RNAGenesis-designed sequences in CRISPR-Cas9 (GFP, B2M, AAVS1), Prime Editing, and Base Editing contexts. These results verify the utility of RNAGenesis in therapeutic genome editing applications.

### Performance of RNAGenesis on BEACON Benchmark

To evaluate the versatility and effectiveness of RNAGenesis, we adopted the BEACON benchmark^15^, which spans three foundational categories—Structure Prediction, RNA Function Prediction, and Engineering Prediction—that are critical for applications such as 3D structural modeling, aptamer design, and CRISPR-based RNA engineering.

As shown in Table 1, RNAGenesis achieves state-of-the-art performance on 10 out of 13 downstream tasks, outperforming recent RNA foundation models, including RNA-FM^6^, RNABERT^5^, and UTRBERT variants^9^. For example, RNAGenesis achieves an *R*^2^ score of 89.03 on alternative polyadenylation site prediction (APA), 97.82% accuracy on non-coding RNA classification (NcRNA), and an *R*^2^ of 85.83 on mean ribosome loading (MRL), significantly outperforming all baselines. These results reflect the model’s superior capability in capturing both regulatory and translational features of RNA, which are crucial for understanding gene expression control and post-transcriptional regulation. As for the secondary structure prediction task, we further benchmarked with RiNALMo^10^ and AIDO.RNA^11^, the latest and largest RNA Foundation models so far in Fig. S5. RNAGenesis also achieves comparable or better performance on Precision, Recall, and F1 values, showing the strong capability of RNAGenesis for structure prediction. In Supplementary, we further perform ablation studies as well as scaling trend analysis of RNAGenesis. Together, these strong results across diverse tasks highlight the broad applicability and advanced modeling capacity of RNAGenesis for RNA biology and therapeutics.

**Table 1.** Benchmark results of 13 RNA tasks from BEACON benchmark. Three color scales of pink first and second are used to denote best performance among all the methods. The baseline results are borrowed from BEACON. ^15^**. Mean (%) is reported for each experiment.**

| Task | SSP | CMP | DMP | SSI | SPL | APA | NcRNA | Modif | MRL | VDP | PRS | CRI-On | CRI-Off |
| --- | --- | --- | --- | --- | --- | --- | --- | --- | --- | --- | --- | --- | --- |
| Metric | F1 (%) | P@L (%) | $R^2$ (%) | $R^2$ (%) | ACC@K (%) | $R^2$ (%) | ACC (%) | AUC (%) | $R^2$ (%) | MCRMSE↓ | $R^2$ (%) | SC (%) | SC (%) |
| CNN | 49.95<sub>(0.82)</sub> | 43.89<sub>(5.53)</sub> | 27.76<sub>(5.00)</sub> | 34.36<sub>(0.12)</sub> | 8.43<sub>(0.38)</sub> | 50.93<sub>(0.17)</sub> | 88.62<sub>(0.71)</sub> | 70.87<sub>(0.4)</sub> | 74.13<sub>(0.58)</sub> | 0.361<sub>(0.003)</sub> | 45.40<sub>(0.66)</sub> | 29.69<sub>(2.52)</sub> | 11.40<sub>(0.10)</sub> |
| ResNet | 57.26<sub>(3.14)</sub> | 59.59<sub>(0.68)</sub> | 30.26<sub>(1.81)</sub> | 37.74<sub>(0.16)</sub> | 21.15<sub>(1.56)</sub> | 56.45<sub>(0.94)</sub> | 88.33<sub>(1.22)</sub> | 71.03<sub>(0.32)</sub> | 74.34<sub>(0.22)</sub> | 0.349<sub>(0.003)</sub> | 55.21<sub>(0.28)</sub> | 28.55<sub>(2.42)</sub> | 11.50<sub>(0.22)</sub> |
| LSTM | 58.61<sub>(0.21)</sub> | 40.41<sub>(1.67)</sub> | 44.77<sub>(0.47)</sub> | 35.44<sub>(1.13)</sub> | 36.66<sub>(1.83)</sub> | 67.03<sub>(0.86)</sub> | 88.78<sub>(0.10)</sub> | 94.83<sub>(0.31)</sub> | 83.94<sub>(0.08)</sub> | 0.329<sub>(0.002)</sub> | 55.45<sub>(0.71)</sub> | 26.83<sub>(1.32)</sub> | 8.60<sub>(0.13)</sub> |
| RNA-FM | 68.50<sub>(0.54)</sub> | 47.56<sub>(6.73)</sub> | 51.45<sub>(0.51)</sub> | 42.36<sub>(0.24)</sub> | 34.84<sub>(0.87)</sub> | 70.32<sub>(0.97)</sub> | 96.81<sub>(0.061)</sub> | 94.98<sub>(0.042)</sub> | 79.47<sub>(0.47)</sub> | 0.347<sub>(0.003)</sub> | 55.98<sub>(0.09)</sub> | 31.62<sub>(1.16)</sub> | 2.49<sub>(1.56)</sub> |
| RNABERT | 57.27<sub>(0.30)</sub> | 45.21<sub>(10.87)</sub> | 48.19<sub>(0.64)</sub> | 31.62<sub>(0.64)</sub> | 0.18<sub>(0.18)</sub> | 57.66<sub>(2.11)</sub> | 68.95<sub>(7.285)</sub> | 82.82<sub>(19.09)</sub> | 29.79<sub>(20.15)</sub> | 0.378<sub>(0.003)</sub> | 54.60<sub>(0.23)</sub> | 29.77<sub>(3.98)</sub> | 4.27<sub>(1.05)</sub> |
| RNA-MSM | 57.98<sub>(0.47)</sub> | 57.26<sub>(15.38)</sub> | 37.49<sub>(4.10)</sub> | 39.22<sub>(0.23)</sub> | 38.33<sub>(0.76)</sub> | 70.40<sub>(1.12)</sub> | 84.85<sub>(0.266)</sub> | 94.89<sub>(0.14)</sub> | 83.48<sub>(0.18)</sub> | 0.330<sub>(0.001)</sub> | 56.94<sub>(0.38)</sub> | 34.92<sub>(1.99)</sub> | 3.85<sub>(0.99)</sub> |
| Splice-H510 | 64.93<sub>(0.84)</sub> | 45.80<sub>(6.03)</sub> | 55.56<sub>(1.00)</sub> | 38.91<sub>(0.07)</sub> | 44.80<sub>(1.93)</sub> | 58.65<sub>(2.34)</sub> | 95.92<sub>(0.666)</sub> | 62.57<sub>(1.92)</sub> | 83.49<sub>(0.47)</sub> | 0.321<sub>(0.000)</sub> | 54.90<sub>(3.45)</sub> | 26.61<sub>(1.30)</sub> | 4.00<sub>(1.13)</sub> |
| Splice-MS510 | 43.24<sub>(28.64)</sub> | 52.64<sub>(7.56)</sub> | 10.27<sub>(0.20)</sub> | 38.58<sub>(0.50)</sub> | 50.55<sub>(0.49)</sub> | 52.46<sub>(17.36)</sub> | 95.87<sub>(0.364)</sub> | 55.87<sub>(5.25)</sub> | 84.98<sub>(0.28)</sub> | 0.315<sub>(0.003)</sub> | 50.98<sub>(7.46)</sub> | 27.13<sub>(0.27)</sub> | 3.49<sub>(2.12)</sub> |
| Splice-MS1024 | 68.26<sub>(0.20)</sub> | 47.32<sub>(3.16)</sub> | 55.89<sub>(0.48)</sub> | 39.22<sub>(0.02)</sub> | 48.52<sub>(0.49)</sub> | 60.03<sub>(3.42)</sub> | 96.05<sub>(0.777)</sub> | 53.45<sub>(6.25)</sub> | 67.15<sub>(30.54)</sub> | 0.313<sub>(0.000)</sub> | 57.72<sub>(0.45)</sub> | 27.59<sub>(4.61)</sub> | 5.00<sub>(0.71)</sub> |
| UTR-LM-MRL | 59.71<sub>(0.30)</sub> | 45.51<sub>(23.51)</sub> | 55.21<sub>(2.91)</sub> | 39.52<sub>(0.36)</sub> | 36.20<sub>(1.84)</sub> | 64.99<sub>(4.90)</sub> | 89.97<sub>(0.617)</sub> | 56.41<sub>(2.90)</sub> | 77.78<sub>(6.03)</sub> | 0.325<sub>(0.002)</sub> | 57.28<sub>(0.10)</sub> | 28.49<sub>(1.37)</sub> | 4.28<sub>(0.15)</sub> |
| UTR-LM-TE&EL | 59.57<sub>(0.20)</sub> | 60.32<sub>(7.27)</sub> | 54.94<sub>(2.54)</sub> | 40.15<sub>(0.11)</sub> | 37.35<sub>(5.48)</sub> | 72.09<sub>(0.82)</sub> | 81.33<sub>(8.551)</sub> | 59.70<sub>(10.52)</sub> | 82.50<sub>(1.45)</sub> | 0.319<sub>(0.001)</sub> | 53.37<sub>(3.54)</sub> | 32.49<sub>(4.14)</sub> | 2.91<sub>(1.18)</sub> |
| UTRBERT-3mer | 60.37<sub>(0.47)</sub> | 51.03<sub>(21.48)</sub> | 50.95<sub>(0.44)</sub> | 34.31<sub>(0.00)</sub> | 44.24<sub>(0.53)</sub> | 69.52<sub>(4.56)</sub> | 92.88<sub>(0.379)</sub> | 95.14<sub>(0.11)</sub> | 83.89<sub>(0.13)</sub> | 0.337<sub>(0.002)</sub> | 56.83<sub>(0.26)</sub> | 29.92<sub>(1.95)</sub> | 4.48<sub>(1.12)</sub> |
| UTRBERT-4mer | 59.41<sub>(0.45)</sub> | 44.91<sub>(27.56)</sub> | 47.77<sub>(2.08)</sub> | 33.22<sub>(0.00)</sub> | 42.04<sub>(0.53)</sub> | 72.71<sub>(0.85)</sub> | 94.32<sub>(0.946)</sub> | 95.10<sub>(0.12)</sub> | 82.90<sub>(0.75)</sub> | 0.341<sub>(0.002)</sub> | 56.43<sub>(0.67)</sub> | 23.20<sub>(1.10)</sub> | 3.11<sub>(1.10)</sub> |
| UTRBERT-5mer | 47.92<sub>(8.75)</sub> | 44.71<sub>(7.64)</sub> | 48.67<sub>(1.70)</sub> | 31.27<sub>(0.00)</sub> | 39.19<sub>(0.37)</sub> | 72.70<sub>(1.77)</sub> | 93.04<sub>(0.367)</sub> | 94.78<sub>(0.07)</sub> | 75.64<sub>(4.70)</sub> | 0.343<sub>(0.001)</sub> | 57.16<sub>(0.08)</sub> | 25.74<sub>(0.00)</sub> | 3.93<sub>(0.24)</sub> |
| UTRBERT-6mer | 38.56<sub>(28.76)</sub> | 51.56<sub>(20.30)</sub> | 50.02<sub>(1.05)</sub> | 29.93<sub>(0.17)</sub> | 38.58<sub>(2.72)</sub> | 71.17<sub>(2.30)</sub> | 93.12<sub>(0.168)</sub> | 95.08<sub>(0.17)</sub> | 83.60<sub>(0.39)</sub> | 0.340<sub>(0.001)</sub> | 57.14<sub>(0.12)</sub> | 28.60<sub>(1.55)</sub> | 4.90<sub>(0.57)</sub> |
| BEACON-B | 64.18<sub>(0.44)</sub> | 60.81<sub>(1.70)</sub> | 56.28<sub>(0.41)</sub> | 38.78<sub>(0.18)</sub> | 37.43<sub>(1.43)</sub> | 70.59<sub>(0.91)</sub> | 94.63<sub>(0.16)</sub> | 94.74<sub>(0.20)</sub> | 72.29<sub>(0.28)</sub> | 0.320<sub>(0.001)</sub> | 54.67<sub>(0.36)</sub> | 26.01<sub>(1.81)</sub> | 4.42<sub>(0.33)</sub> |
| BEACON-B512 | 58.75<sub>(3.72)</sub> | 61.20<sub>(2.11)</sub> | 56.82<sub>(0.63)</sub> | 39.13<sub>(0.08)</sub> | 37.24<sub>(1.09)</sub> | 72.00<sub>(0.17)</sub> | 94.99<sub>(0.21)</sub> | 94.92<sub>(0.07)</sub> | 72.35<sub>(0.28)</sub> | 0.320<sub>(0.001)</sub> | 55.20<sub>(0.26)</sub> | 28.17<sub>(1.81)</sub> | 3.82<sub>(1.04)</sub> |
| RNAGenesis | 73.37<sub>(0.25)</sub> | 72.10<sub>(0.23)</sub> | 70.77<sub>(0.33)</sub> | 40.75<sub>(0.08)</sub> | 38.14<sub>(1.39)</sub> | 89.03<sub>(0.02)</sub> | 97.82<sub>(0.02)</sub> | 95.16<sub>(0.10)</sub> | 85.83<sub>(0.05)</sub> | 0.291<sub>(0.001)</sub> | 58.19<sub>(0.11)</sub> | 35.46<sub>(1.23)</sub> | 44.93<sub>(3.77)</sub> |

### RNAGenesis for RNA Structural Tasks

RNA sequences encode rich information about their three-dimensional (3D) structures and functions through evolutionary constraints, offering valuable priors for structural discovery^16, 25^. By integrating RNAGenesis with structure-aware modules, we transform it into a versatile multi-modal framework capable of structure prediction, inverse folding, and *de novo* RNA design (Figure 2a). Specifically, we adopt pretrained structural encoders from RhoDesign and RNA-FrameFlow and fuse them with the pretrained RNAGenesis sequence encoder via a multi-modal integration strategy inspired by^26^. The resulting model is then fine-tuned using LoRA^27^ on diverse structural datasets. This design endows RNAGenesis with rapid adaptability to various structure-guided tasks (e.g., low-resource settings, structure generation), demonstrating strong generalization and scalability under limited supervision.

**Figure 2.**
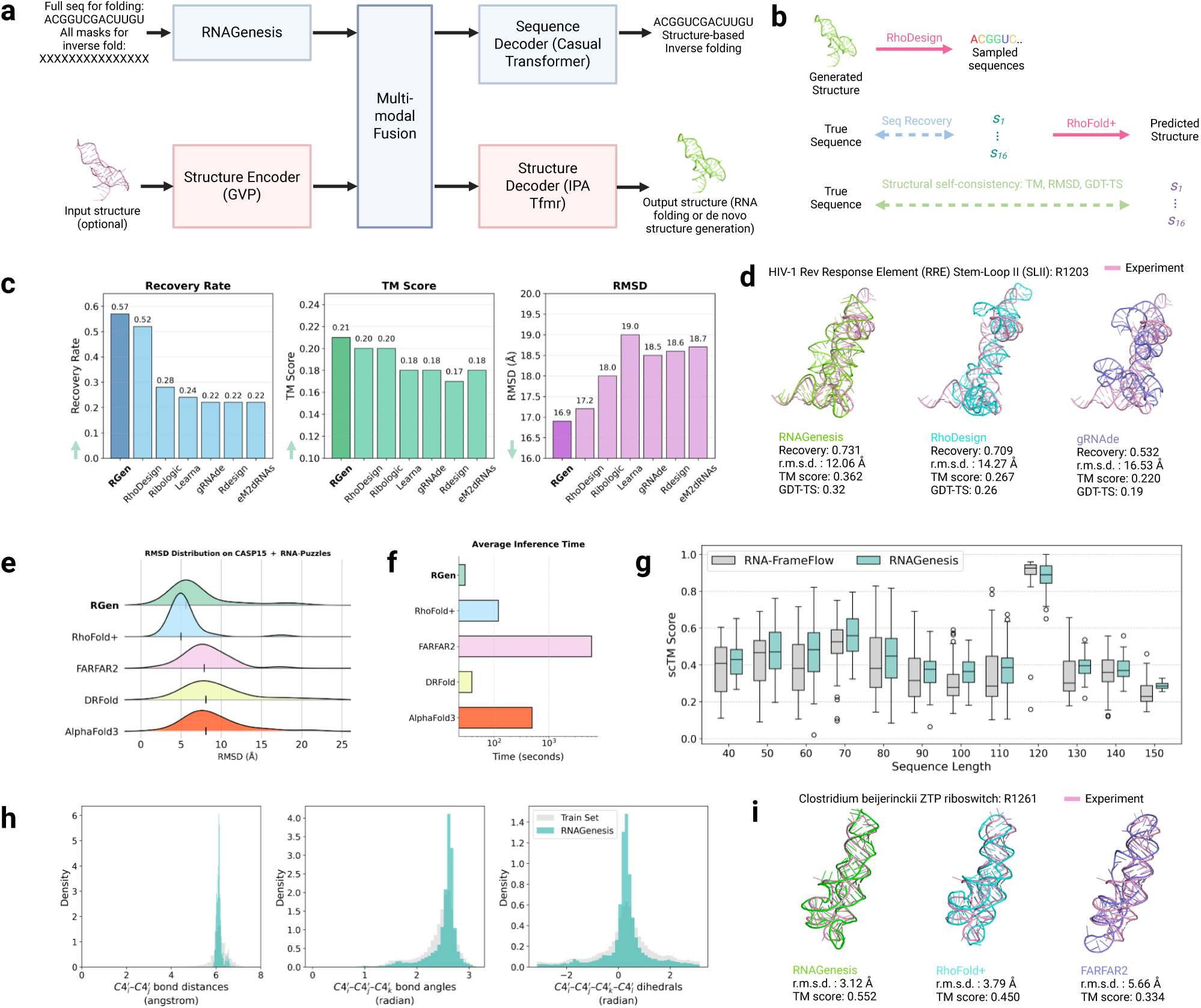
RNAGenesis as a unified model for structural tasks. (**a**) RNAGenesis can be integrated with Structure Encoder/Decoder and Sequence Decoder for multi-modal tasks, including structure-based inverse folding, RNA structure prediction, and *de novo* structure generation. (**b**) In-silico evaluation metrics for structural tasks, including (1) sequence recovery - the percentage of native nucleotides recovered in designed samples, and (2) self-consistency scores - measured by ‘forward folding’ designed sequences using RhoFold+ to assess 3D structure recovery, such as TM score, RMSD, and GDT-TS. (**c**) Performance comparison of RNAGenesis against other models for RNA design on a held-out test set, where higher Recovery Rate and TM Score indicate better performance, while lower RMSD is preferred. (**d**) Case study applying RNAGenesis, RhoDesign, and gRNAde to redesign the HIV-1 Rev Response Element (RRE) Stem-Loop II from CASP16, evaluating sequence recovery, RMSD, TM score, and GDT-TS after inverse folding and refolding with RhoFold+. (**e**) RMSD performance comparison across 6 CASP15 targets and 24 non-overlapping RNA-Puzzles targets, visualized as ridgeline plots. (**f**) Average inference time comparison across different methods. (**g**) Performance comparison between RNAGenesis and RNA-FrameFlow in generating *de novo* RNA structures of varying lengths (40-150 nucleotides), with scTM scores calculated for 50 structures per length. (**h**) Distribution analysis of structural properties from 600 RNAGenesis-generated structures compared to the RNASolo training set, showing inter-nucleotide bond distances, bond angles between nucleotide triplets, and torsion angles between four consecutive nucleotides. (**i**) Case study demonstrating the application of RNAGenesis, RhoFold+, and FARFAR2 in predicting the structure of the Clostridium beijerinckii ZTP riboswitch from CASP16, with RMSD and TM scores reported.

In the **Inverse Folding** setting, the model is given a target RNA tertiary structure and tasked with generating RNA sequences whose folded conformations closely match the input. Building upon RhoDesign^17^, which leverages geometric vector perceptrons (GVPs)^28^ and causal transformers^29, 30^, RNAGenesis extends this framework by fusing structure-conditioned embeddings with its own pretrained sequence encoder^26^. We benchmarked RNAGenesis against leading inverse folding models, including RhoDesign, Ribologic^31^, Learna^32^, gRNAde^33^, RDesign^34^, and eM2dRNAs^35^, using 3,435 experimentally resolved RNA structures from the Protein Data Bank (PDB)^36^ curated by RhoDesign. As shown in Figure 2b, RNAGenesis achieves the best performance across sequence recovery rate, TM-score, and RMSD—achieving a 57.1% recovery rate and 16.9 °A RMSD, outperforming RhoDesign (52.9%, 17.1 °A). To further evaluate structural fidelity, we assess inverse folding on the HIV-1 Rev Response Element (RRE) Stem-Loop II, a well-characterized motif involved in viral replication. Predicted sequences generated by RNAGenesis, RhoDesign, and gRNAde are folded and compared to the CASP16 reference structure using metrics such as RMSD, TM-score, and GDT-TS. As shown in Figure 2d, RNAGenesis consistently produces structures closest to the native fold. Compared with RhoDesign and gRNAde, it achieves the best trade-off between sequence diversity and structural accuracy, demonstrating strong controllability over the sequence–structure relationship.

**RNA three-dimensional (3D) structure prediction** is a long-standing and unresolved challenge in molecular biology^37^. However, the inherent structural flexibility of RNA, combined with the scarcity of high-resolution experimental data, makes computational prediction particularly difficult. By combining the RNAGenesis sequence encoder with a structure decoder (specifically, the IPA transformer from RNA-FrameFlow^24^) and fine-tuning on the RNASolo dataset^38^, RNAGenesis gains the ability to predict RNA 3D structures directly from sequence. Following the evaluation framework of RhoFold+^39^, we benchmarked RNAGenesis against state-of-the-art RNA structure prediction models, including RhoFold+^39^, FARFAR2^40^, DRFold^41^, and AlphaFold3^20^, on two rigorous datasets: CASP15 RNA targets^32^ and the RNA-Puzzles benchmark^42^, which are widely recognized for evaluating RNA folding accuracy on challenging and diverse structures. As illustrated in Figure 2e,f, RNAGenesis achieves the second-best performance in terms of root-mean-square deviation (RMSD), closely matching RhoFold+ and significantly outperforming all other baselines. In addition, RNAGenesis demonstrates the *fastest inference speed* among all compared methods, thanks to its streamlined sequence-to-structure architecture that avoids expensive atomistic simulations and multiple sequence alignment (MSA) searches. These results underscore the potential of RNAGenesis as a strong foundation model for RNA structure prediction—offering competitive accuracy with state-of-the-art methods, while maintaining superior generalization ability and runtime efficiency due to its simple yet effective design. To further highlight its structural modeling capabilities, we present a case study in Figure 2i using the Clostridium beijerinckii ZTP riboswitch, a metabolite-sensing non-coding RNA element that regulates gene expression in response to cellular ZTP levels. We compare RNAGenesis, RhoFold+, and FARFAR2 in terms of both RMSD and TM-score, measuring atomic-level precision and global structural similarity, respectively. RNAGenesis achieves the best accuracy with an RMSD of 3.12 °A versus 3.79 °A for RhoFold+, and a TM-score of 0.552 compared to 0.450. These results show that RNAGenesis can more accurately capture both local and global structural features, even on complex functional RNAs such as riboswitches. In future work, we plan to incorporate MSAs into RNAGenesis to further enhance prediction accuracy, as MSA-derived features have been shown to improve RNA structure modeling in recent studies^39^.

Finally, we adapt RNAGenesis for the task of ***de novo* RNA structure generation**, where the model generates RNA 3D structures without any conditioning sequence or structural template. To enable this, we fully mask the input tokens to the sequence encoder, effectively removing sequence-specific information. Notably, although RNAGenesis uses the same decoder architecture as RNA-FrameFlow^24^, it consistently produces higher-quality structures. This improvement can be attributed to RNAGenesis’s pretrained encoder, which has been trained on large-scale sequence-to-structure supervision and learns a biologically meaningful latent space. Even under full masking, this encoder generates informative representations that guide the decoder toward physically plausible and well-folded RNA structures—a capability lacking in RNA-FrameFlow, which is trained purely in the unconditional setting. As shown in Figure 2g, we evaluate *de novo* RNA structure generation across a range of sequence lengths (40–150 nucleotides), generating 50 structures per length and assessing global structural quality using the scTM score. RNAGenesis consistently outperforms RNA-FrameFlow across all length ranges, achieving higher scTM scores and indicating superior recovery of native-like global folds. This demonstrates the model’s robustness in handling both short and long RNAs, reflecting strong cross-scale generalization capabilities. Furthermore, analysis of 600 generated structures reveals that RNAGenesis also preserves realistic local geometry—bond lengths and angles that closely match experimentally resolved RNA structures (Figure 2h). These results highlight RNAGenesis’s capacity to jointly model global fold fidelity and local stereochemical accuracy in *de novo* RNA structure generation.

### RNAGenesis Applied for Diverse RNA Therapeutic Modalities

RNA therapeutics have emerged as a transformative class of medicines, fundamentally altering the approach to disease treatment by utilizing RNA molecules to interact directly with cellular nucleic acid or to modulate protein production^43^. In Figure 3, we introduce **RNATx-Bench**, a benchmark we constructed to systematically evaluate RNA foundation models on RNA therapeutics. It comprises over 100,000 experimental and clinical datapoints spanning five major therapeutic modalities: antisense oligonucleotides (ASOs), circular RNAs (circRNAs), RNA aptamers, small interfering RNAs (siRNAs), and short hairpin RNAs (shRNAs). Additional details are provided in the Supplementary Information.

**Figure 3.**
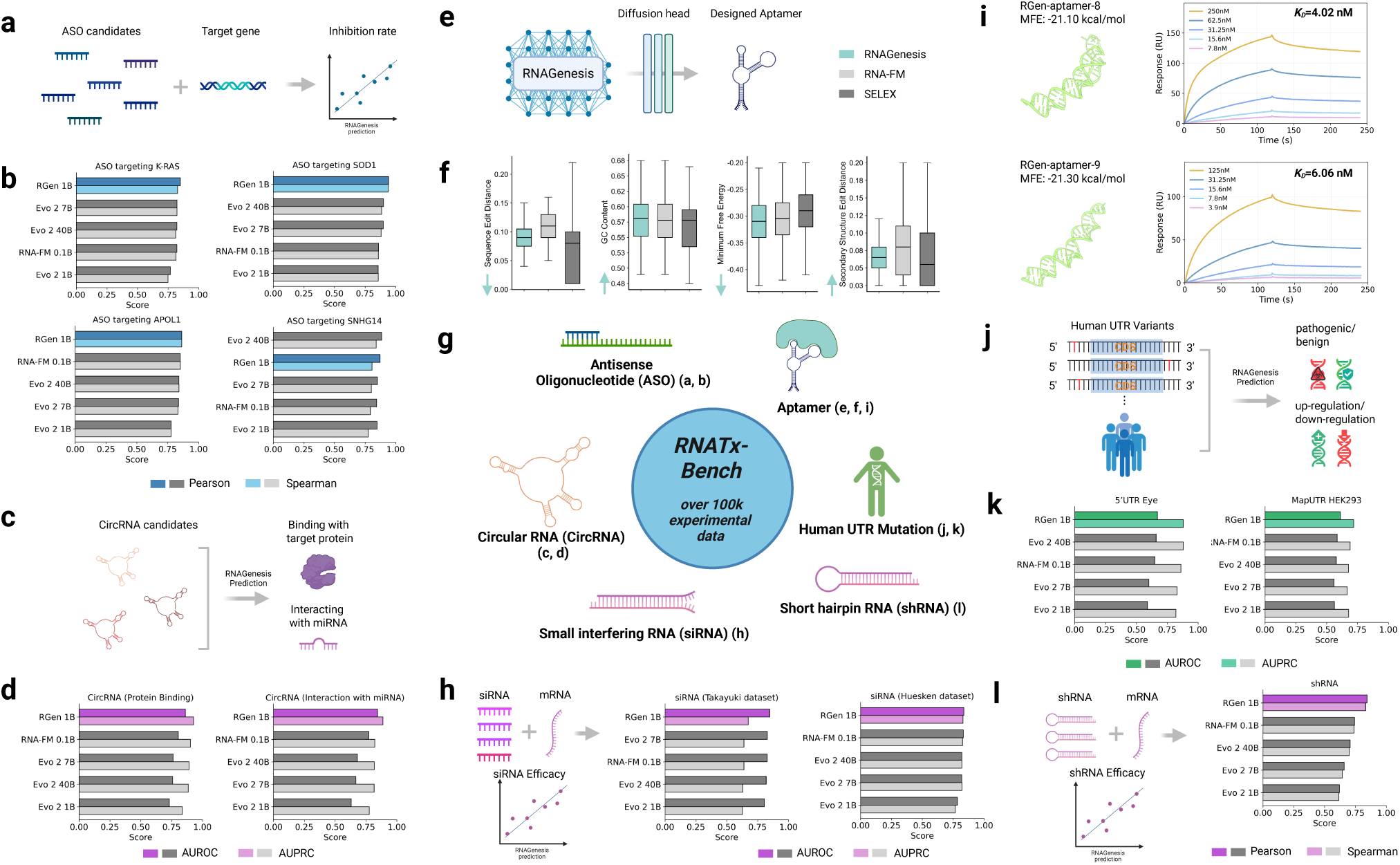
RNAGenesis accurately predicts the efficacy of diverse RNA therapeutics and the functional impact of human UTR variants on RNATx-Bench. **a**, **b**, Performance of RNAGenesis and baseline models in predicting antisense oligonucleotide (ASO) inhibition rates for four representative target genes (*N* = 943, 2,264, 2,140, and 1,890 for K-RAS, SOD1, APOL1, and SNHG14, respectively. Total *N* = 34,773 for ASO dataset). **c**, **d**, Prediction of circular RNA (circRNA) functionality, including classification of binding to target proteins (**c**; *N* = 2,072) and interaction with microRNAs (**d**; *N* = 2,092). **e**, Schematic of the *de novo* aptamer design workflow using latent diffusion with RNAGenesis. **f**, Comparison of key biophysical properties for aptamer sequences (sequence edit distance, G/C content, minimum free energy, and secondary structure edit distance). Distributions are shown for sequences generated by RNAGenesis (25,600 sequences), a baseline model (RNA-FM; 25,600 sequences), and experimentally-derived sequences from HT-SELEX (13,802 sequences). **g**, Overview of RNA therapeutic modalities and variant types included in RNATx-Bench, a unified benchmark we constructed comprising over 100,000 experimentally validated datapoints across ASO, siRNA, shRNA, aptamer, circRNA, and human UTR variant assays. **h**, Performance on predicting small interfering RNA (siRNA) efficacy using the Takayuki (*N* = 702) and Huesken (*N* = 2,361) datasets. **i**, AlphaFold3-predicted tertiary structures of a representative aptamers designed by RNAGenesis with higher binding affinity than wild type. For each structure, the minimum free energy (MFE) and experimentally measured dissociation constant (*K*_*D*_) from surface plasmon resonance SPR assays are reported. **j**, **k**, Performance on predicting the effects of human UTR variants. Tasks include classifying variants as pathogenic or benign (**j**; *N* = 1,547 for 5’UTR Eye dataset) and as up-regulating or down-regulating (**k**; *N* = 17,210 for MapUTR HEK293 dataset). **l**, Performance on predicting short hairpin RNA (shRNA) efficacy (*N* = 2,076). In **f**, box plots show the median (centre line), interquartile range (IQR; box bounds), and whiskers extending to 1.5× the IQR. In **b**, **d**, **h**, **k**, and **l**, RGen: RNAGenesis.

**Antisense Oligonucleotides (ASOs)** are short, synthetic, single-stranded nucleic acid molecules, offering a promising treatment for a wide range of genetic and rare diseases^44^. However, due to the huge search space and varying molecular mechanisms, optimizing and screening ASO candidates is a challenging task^45^. In Figure. 3 a & b, we collect ASO candidates and their experimental inhibition rates targeting 4 clinically critical genes, including **SOD1**, **K-RAS**, **APOL1**, and **SNHG14**. In Figure. 3, RNAGenesis achieves the highest Pearson/Spearman correlation scores in 3 out of 4 tasks. The high correlation between RNAGenesis predictions and experimental inhibition data for these targets underscores the model’s utility in guiding ASO design and prioritization.

**Circular RNAs (circRNAs)** represent a distinct and increasingly recognized class of single-stranded RNA molecules characterized by a covalently closed loop structure^46^. Due to their circular conformation and lack of free ends, circRNAs exhibit enhanced stability and play critical roles in gene regulation and the pathogenesis of various complex diseases. Accurate identification and prediction of circRNA regulatory interactions are essential for understanding their biogenesis and functional impact. In Figure 3, we evaluate the ability of RNA foundation models to predict regulatory features of circRNAs, including their interactions with microRNAs and RNA-binding proteins. Compared to existing models such as RNA-FM and Evo2, **RNAGenesis** demonstrates a clear advantage. For instance, RNAGenesis achieves an AUROC of 0.8619, outperforming RNA-FM (0.8047) and Evo2-40B (0.7576). This indicates the superior capability of RNAGenesis in modeling circRNA regulatory landscapes, highlighting its potential for advancing RNA-based therapeutics.

**Small interfering RNAs (siRNAs)** are short double-stranded RNA molecules (20–25 base pairs) that mediate gene silencing via the conserved RNA interference (RNAi) pathway^47^. By guiding the RNA-induced silencing complex (RISC) to degrade target mRNAs, siRNAs can selectively silence disease-related genes, making them a powerful modality for therapeutic intervention. However, designing effective siRNAs remains challenging, requiring high target specificity, minimal immune activation, and low off-target toxicity. In Figure 3 h, we demonstrate the application of RNAGenesis for predicting siRNA efficacy, achieving an AUROC exceeding 0.8—surpassing all other RNA/DNA foundation models. This highlights RNAGenesis’s strong potential for guiding siRNA design in therapeutic settings.

**Short hairpin RNAs (shRNAs)** are single-stranded RNA molecules that can mediate gene silencing to achieve RNA interference therapeutics^48^. While siRNAs are easy to synthesize and widely used for gene knockdown, their effects typically last only a few days in cells, limiting their utility in both research and therapeutic contexts. In contrast, short hairpin RNAs (shRNAs) can achieve similarly effective gene silencing while enabling more sustained knockdown, thereby expanding their potential for long-term studies and clinical applications^49^. In Figure 3 l, we apply RNAGenesis to predict the efficacy of shRNA candidates, achieving a notable improvement in prediction correlation—over 10% higher than baseline methods—demonstrating its superior capability in modeling shRNA functionality.

**Aptamers** are short, single-stranded RNA molecules capable of folding into complex 3D structures that bind specific targets (e.g., protein) with high affinity and specificity. Their programmable nature makes them powerful tools in therapeutics, diagnostics, and synthetic biology^50–53^. To assess the utility of RNAGenesis for de novo aptamer design, we evaluated its ability to generate sequences that match the key characteristics of real aptamers derived from the HT-SELEX dataset A, provided by AptaDiff^54^. SELEX (Systematic Evolution of Ligands by EXponential enrichment) is a widely adopted method for identifying high-affinity RNA binders. Dataset A contains 13,802 RNA sequences enriched through six rounds of SELEX targeting IGFBP3, a biologically relevant protein involved in growth regulation and cancer progression^55^. These high-quality aptamer candidates serve as the benchmark for evaluating generative models.

We adapted RNAGenesis by training a latent diffusion model with a frozen encoder to generate aptamer sequences in the learned embedding space. As shown in Figure 3 f, we compared 25,600 sequences generated by RNAGenesis and RNA-FM (batch size 128, resampled 200 times) against the SELEX dataset using four key evaluation metrics: sequence edit (Levenshtein) distance, GC content, minimum free energy (MFE), and RNA secondary structure edit distance. Across all metrics, RNAGenesis demonstrates superior performance: it produces sequences with smaller sequence and secondary structure edit distances, lower minimum free energy (MFE) values, and higher GC content, indicating a stronger alignment with the characteristics of natural SELEX-derived aptamers. Lower MFE values reflect greater thermodynamic stability, suggesting that the generated sequences are more likely to fold into stable, well-defined secondary structures. Meanwhile, higher GC content is associated with stronger base-pairing interactions, due to the triple hydrogen bonds in G-C pairs, further contributing to structural robustness and folding reliability. Additionally, Figure 3 i presents AlphaFold3-predicted 3D structures of a SELEX-derived aptamer and two top-performing aptamers designed by RNAGenesis, annotated with their minimum free energy (MFE) values and experimentally measured dissociation constants (*K*_*D*_) from SPR assays. Both RGen-aptamer-8 and RGen-aptamer-9 exhibit stronger binding affinities to the target protein compared to the SELEX control, with *K*_*D*_ values of 4.02 nM and 6.06 nM, respectively, versus 11.6 nM for the SELEX aptamer (Figure. S15, S16, and S17). In addition, the RNAGenesis-designed aptamers display slightly lower MFE values, suggesting increased thermodynamic stability. These results demonstrate that RNAGenesis generates aptamers with enhanced binding affinity and favorable folding energetics, validating its utility in functional RNA design.

### RNAGenesis for Human UTR Variants Effect Prediction

Here, we evaluate the performance of RNAGenesis in predicting the functional consequences of human UTR variants, first on clinical datasets for pathogenicity and then on large-scale experimental datasets for regulatory effects. The 5’ and 3’ untranslated regions (UTRs) of messenger RNA play a critical role in post-transcriptional regulation, yet variants within these regions remain a major interpretive challenge in genomic medicine. Such variants are an often overlooked source of Mendelian diseases, including inherited retinal diseases (IRDs)^56^, but predicting their functional effects computationally is notoriously difficult^57^. To address this challenge, we evaluated the ability of our model, RNAGenesis, alongside baseline RNA/DNA foundation models to predict the pathogenicity of UTR variants. Using comprehensive variant datasets for IRDs^56^ and from ClinVar^57^, we demonstrate that RNAGenesis achieves state-of-the-art performance, ranking either first or second among the models tested (Fig. 3 j). This high accuracy in distinguishing pathogenic from benign variants underscores the model’s potential as a powerful tool for clinical variant prioritization and for improving the diagnostic yield in genetic diseases.

Furthermore, to assess the model’s ability to predict more nuanced regulatory activity, we benchmarked it using data from massively parallel reporter assays (MPRAs). These experiments have generated invaluable datasets, such as Maptur^58^, that quantify the impact of thousands of variants on mRNA abundance, typically measured as a log-fold change (lnfc). For this task, we trained RNAGenesis and baseline models to classify variants as significantly down-regulating, up-regulating, or neutral based on their experimental lnfc values in HEK293 and HELA cell lines. Successfully classifying a variant’s regulatory consequence is critically important, as it enables the high-throughput prioritization of functional variants for disease studies and deepens our understanding of the sequence-based rules governing gene expression. As observed in Figure 3k and Figure. S13, RNAGenesis significantly outperforms all other baselines, indicating its superior ability to decipher the complex sequence determinants that dictate a variant’s regulatory impact.

### RNAGenesis-designed sgRNA Scaffold Enhances CRISPR-Cas9 Editing Efficiency

The CRISPR/Cas system has revolutionized genome engineering by enabling precise DNA targeting through RNA-guided endonucleases, where the single guide RNA (sgRNA) directs the Cas9 protein to induce double-stranded breaks (DSBs) for gene editing^4, 59–62^. The sgRNA consists of a 20-nucleotide spacer sequence complementary to the target DNA and a scaffold domain that binds Cas9, but its efficacy is often limited by intramolecular interactions leading to undesirable secondary structures, which restrict editing efficiency at many genomic loci^63^. To address this, advances in CRISPR technology, such as Cas9 variants (e.g., nCas9, dCas9) and sgRNA modifications, have expanded their applications, yet sequence inflexibility remains a challenge. In RNAGenesis, we combine strong generation capabilities and bio insights to design sgRNAs with enhanced properties (Fig. 4 a). We trained a classifier for gradient guidance^64^ using the 154 DNA cleavage in vitro data from^63^, enabling precise steering of the denoising process toward high-cleavage-efficiency sequences (more details of the data are included in Supplementary). For beam search optimization, we incorporated secondary structure prediction and minimum free energy (MFE) calculations using RNAFold and ViennaRNA^65^, defining the reward as a combination of structural similarity (ss similarity) and the ratio of predicted MFE to wild-type MFE. This reward function guides the selection of 1 beam expanded into 8 candidates per timestep (as per Algorithm 1), iteratively refining latents to maximize editing potential. Finally, the designed sgRNA sequences are ranked using a classifier trained on the 5-round Binding and Ligand-Activated Directed Evolution (BLADE) SELEX data^63^, ensuring the selection of top-performing variants. This integrated approach leverages RNAGenesis’s generative capabilities, combining gradient guidance’s precision with beam search’s diversity, to produce sgRNAs that address the limitations of traditional CRISPR targeting, achieving enhanced cleavage efficiency and structural stability through inference-time alignment. By generating a large pool of sequences and applying a series of stringent filters for minimum free energy (MFE), secondary structure similarity, and homology, RNAGenesis consistently yielded a greater number of high-quality candidate sequences compared to other generative models (Fig. 4 b & c). We then tested the top-ranked scaffolds experimentally. In a GFP reporter knockout assay, multiple RNAGenesis-designed scaffolds outperformed the conventional wild-type (WT) guide, with the top candidate increasing editing efficiency by 13% (Fig. 4 d). Overall, 6 of the 24 tested designs showed significantly higher activity (Fig. S8). This enhanced performance was confirmed at endogenous loci, where the RNAGenesis-6 (RGen-6) scaffold achieved more effective knockout of both the B2M (Fig. S12) and AAVS1 (Fig. 4 d) genes across a range of sgRNA dosages (e.g., around 2-fold efficiency improvement on AAVS1 medium dosage). These results establish that our design strategy successfully generates novel sgRNA scaffolds that increase the editing efficiency of the standard CRISPR-Cas9 system.

**Figure 4.**
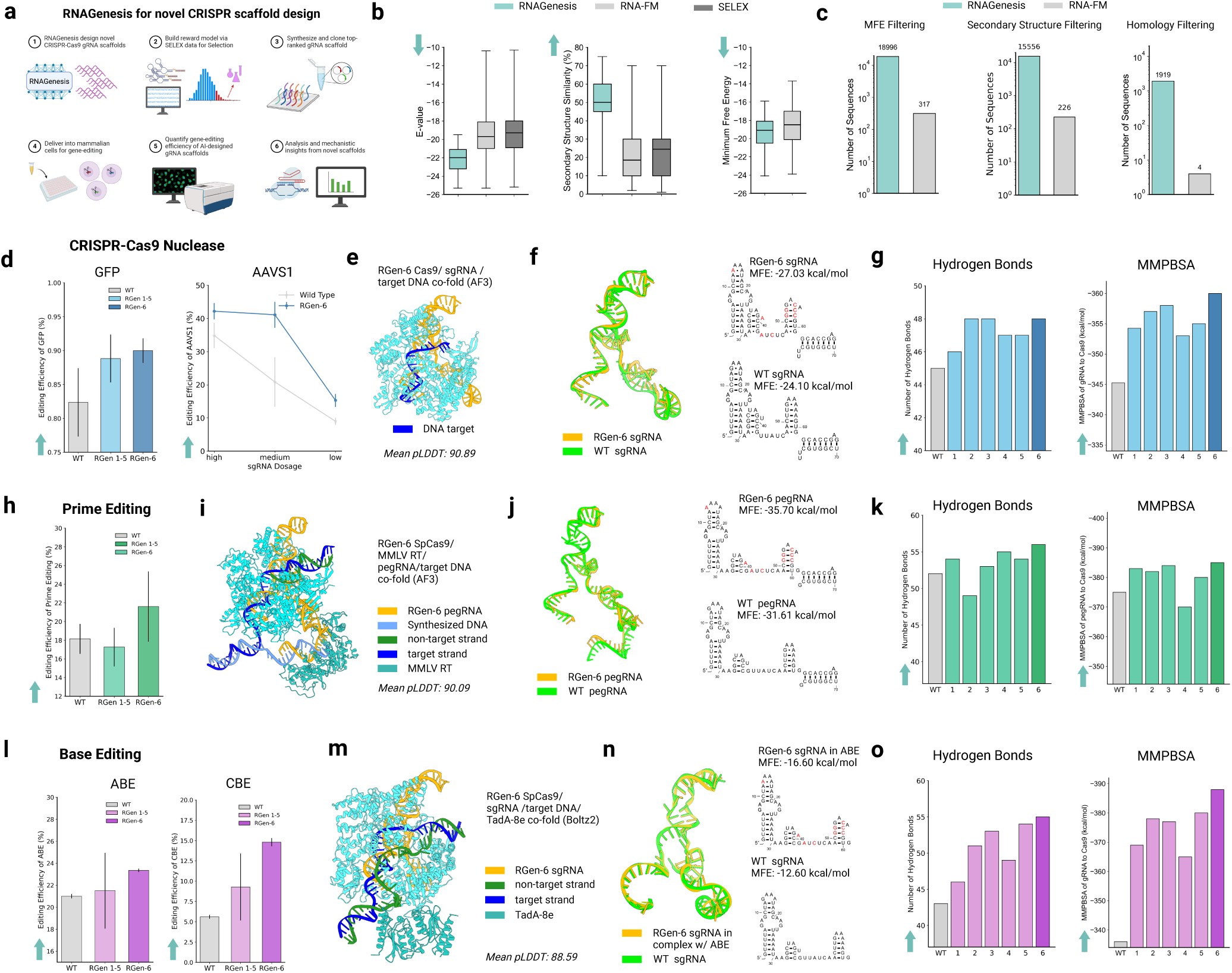
The RNAGenesis-designed sgRNA scaffold enhances the efficiency of CRISPR-Cas9, prime editing, and base editing systems through improved structural stability. (**a**) Workflow for RNAGenesis-designed CRISPR-Cas9 sgRNA scaffolds: design of novel scaffolds, reward model-based selection using SELEX data, synthesis and cloning of top-ranked scaffolds, delivery into mammalian cells, quantification of gene-editing efficiency, and mechanistic analysis with computational tools. (**b**) Comparison of the E-value, secondary structure similarity, and minimum free energy of the generated sgRNA sequences. For RNAGenesis and RNA-FM, 25600 sequences are generated (batch size = 128 and sampling 200 times) with the same latent diffusion architecture. The SELEX dataset is shown as a reference. (**c**) Filtering sgRNA sequences based on MFE (absolute difference less than 2 with wild type), secondary structure similarity (greater than 60%), and high homology (1e-28 ≤ e-value ≤ 1e-25). (**d**) Comparative analysis of CRISPR-Cas9 editing efficiency for sgRNAs targeting GFP and AAVS1. For GFP, both the average of five RNAGenesis scaffolds (RGen 1-5) and a single top guide (RGen-6) were compared with the wild-type (WT) control. The RNAGenesis RGen-6 guide, in particular, demonstrates superior performance for AAVS1 across various plasmid dosages (high: 400ng, medium: 250ng, low: 100ng). (**e**) AlphaFold3 (AF3) structure prediction of RGen-6 sgRNA in complex with Cas9 and target DNA. (**f**) RGen-6 sgRNA aligned to the wild-type sgRNA from PDB 4OO8. Secondary structure comparison between RGen-6 and wild-type sgRNA, with mutations highlighted in red. MFE is calculated with RNAFold. (**g**) Computational analysis of sgRNA-Cas9 interactions. Bar charts show the number of hydrogen bonds between and the calculated MMPBSA binding energy for the RNAGenesis scaffolds compared to WT. (**h**) Prime Editing efficiency using RNAGenesis-designed pegRNAs. (**i**) AlphaFold3 (AF3) structure prediction of RGen-6 prime editing guide RNA (pegRNA) in complex with SpCas9, MMLV Reverse Transcriptase, and target DNA. (**j**) RGen-6 pegRNA aligned to the wild type pegRNA from PDB 6WUT. Secondary structure comparison between RGen-6 and wild-type pegRNA, with mutations highlighted in red. (**k**) Computational analysis of pegRNA-Cas9 interactions. Bar charts show the number of hydrogen bonds between and the calculated MMPBSA binding energy for the RNAGenesis scaffolds compared to WT. (**l**) Base Editing efficiency using RNAGenesis-designed sgRNAs for both Adenine (ABE) and Cytosine (CBE) base editors. (**m**) Structural prediction (Boltz2) of an Adenine Base Editor, showing the SpCas9 protein fused to the TadA-8e deaminase and guided to its target DNA by the RGen-6 sgRNA. (**n**) RGen-6 sgRNA in complex with ABE aligned to the wild-type sgRNA from PDB 6VPC. The secondary structure comparison is shown on the right. (**o**) Computational analysis of sgRNA-Cas9 interactions in Adenine Base Editor. Bar charts show the number of hydrogen bonds and the calculated MMPBSA binding energy for the RNAGenesis scaffolds compared to WT.

### RNAGenesis Scaffold Transfers to Base Editing and Prime Editing

Base editing^18^ and prime editing^19^ represent major advances in gene editing, enabling precise single-base substitutions as well as small insertions or deletions without inducing potentially deleterious double-strand breaks. These technologies hold immense therapeutic potential, as a large fraction of known human pathogenic genetic variants are point mutations that could theoretically be corrected by such tools. However, the clinical translation of these advanced editors is often hindered by their variable and sometimes insufficient efficiency. We therefore sought to determine if our RNAGenesis-designed scaffolds could enhance these therapeutically relevant systems. A key test of our model’s ability to capture fundamental design principles was its capacity for **zero-shot** generalization, as the model was trained exclusively on CRISPR-Cas9 data.

First, we incorporated our designed scaffolds into a prime editing guide RNA (pegRNA). The resulting RNAGenesis pegRNAs successfully improved performance, increasing prime editing efficiency by up to 1.2-fold compared to the wild-type guide (RGen-6 in Fig. 4 h). This enhancement was even more pronounced in base editing systems. For base editing, the RNAGenesis-6 scaffold boosted cytosine base editor (CBE) efficiency by over 2.5-fold compared to the WT scaffold (14.8% vs 5.5%) and also yielded a modest improvement for adenine base editor (ABE) (Fig. 4 l). This zero-shot enhancement underscores RNAGenesis’s ability to capture and transfer fundamental principles of sgRNA structural optimization, offering a versatile and generalizable framework for advancing a wide range of gene editing technologies.

### Computational Analysis Reveals the Structural Basis for Enhanced Activity

To understand the mechanism behind the improved performance, we performed computational analyses. Structural prediction using AlphaFold3^20^ and Boltz-2^21^ generated high-confidence models of the RGen-6/Cas9 complex (mean pLDDT > 90), which aligned closely with the experimental crystal structures (Fig. 4 e, i, m and Fig. S11). The RGen-6 sgRNA was predicted to have a more favorable minimum free energy (−27.03 kcal/mol vs. −24.10 kcal/mol for wild-type), as calculated by RNAFold^65^, suggesting greater thermodynamic stability (Fig. 4 f). Importantly, these mutations preserved the global secondary structure while enhancing local base-pairing stability, notably through an increased frequency of G-C pair formation. Consistent trends were observed in prime editing and base editing contexts. RNAGenesis-designed pegRNAs and sgRNAs retained favorable scaffold conformations and reduced free energy values (e.g., –35.70 vs. –31.61 kcal/mol for pegRNA; –16.60 vs. –12.60 kcal/mol for base editing sgRNA) (Fig. 4 j,n).

Indeed, detailed 3D structural analysis revealed additional hydrogen bonds formed between the RNAGenesis scaffold and key Cas9 residues such as R741 (Fig. S11). Quantitative analyses using MDAnalysis^66^ and Uni-GBSA^67^ demonstrated that RGen-6 consistently forms more hydrogen bonds with Cas9 and exhibits more favorable MMPBSA binding free energies compared to both the wild-type and other RNAGenesis-designed candidates (Fig. 4g,k,o). For example, in the base editing context, RGen-6 forms 56 hydrogen bonds versus 43 for the wild-type, with an MMPBSA binding free energy of –388 kcal/mol compared to –336 kcal/mol for the wild-type scaffold (Fig. 4o). This computationally supported increase in ribonucleoprotein complex stability likely contributes to the enhanced editing efficiencies observed in experiments.

In summary, RNAGenesis designs novel sgRNA scaffolds that significantly enhance editing efficiency across diverse platforms—from CRISPR-Cas9 to therapeutically vital prime and base editors—by improving the scaffold’s structural stability and binding affinity, thus offering a powerful and generalizable approach to advance gene editing applications.

## Discussion

RNAGenesis represents a major advance in bridging RNA sequence modeling with de novo structural and functional design. By unifying latent diffusion, structural priors, and inference-time alignment, it enables programmable synthesis of functional RNAs across a wide range of therapeutic and genome editing tasks. Extensive evaluations on the BEACON and RNATx-Bench benchmarks—coupled with strong experimental validation across aptamer design and gene editing modalities, including CRISPR, base editing, and prime editing—underscore RNAGenesis as a versatile and generalist foundation model for RNA biology and therapeutics.

Several key directions remain for future exploration. A critical next step is the development of a unified biomolecular model capable of jointly explicitly modeling DNA, RNA, and proteins—thus capturing the full flow of genetic information across the central dogma^68^. Such a model would enable integrated understanding of transcriptional regulation, RNA processing, and translation, while supporting multimodal tasks such as codon optimization, RNA–protein binding prediction, and regulatory element design. Within this broader framework, it remains essential to accurately capture the distinct sequence organization, structural constraints, and functional logic of both coding and non-coding RNAs. Incorporating these distinctions into a unified system could improve generalization, reduce reliance on task-specific finetuning, and expand applicability across molecular contexts. Furthermore, scaling experimental validation across diverse cell types, therapeutic modalities, and delivery systems will be key to fully realizing the translational potential of foundation models in RNA biology and beyond.

Moreover, integrating RNAGenesis with autonomous AI agents offers a powerful paradigm for closed-loop biological design. Recent advances in scientific agents capable of hypothesis generation, simulation orchestration, and experimental planning open the door to agent-driven RNA therapeutics pipelines. By combining RNAGenesis with such agents, one could enable continuous cycles of design, evaluation, and optimization—accelerating drug discovery, synthetic biology, and the realization of in silico cell systems. In this broader context, RNAGenesis serves not only as a foundation model but as a foundation for programmable biology.

## Methods

### Model Architecture

#### Hybrid N-Gram Tokenization

A hybrid tokenization scheme is adopted including single nucleotide tokenization and N-Gram tokenization with CNN. RNA sequences are primarily composed of four nucleotide units A,U,C,G, and single nucleotide tokenization can represent the basic information. The N-Gram tokenization based on 1D CNN with different Kernel sizes can learn the higher level of interactions such as RNA motifs. The CNN transformation is unique in RNAGenesis compared with baseline models such as RiNALMo^10^ and Uni-RNA^11^. The input into the Encoder is the sum of the hybrid tokenization representations. After tokenization, each nucleotide is transformed into a 1280-dimensional vector.

#### Encoder

The backbone of RNAGenesis is an Bert-style encoder-only Transformer that focuses on RNA sequence modeling and understanding. We denote the encoder by Enc which maps a sequence (*x* = *x*^1^, *x*^2^,…, *x*^*L*^) to a sequence of contextualized token embeddings of the same length *L*. At its core lies the self-attention mechanism^29^, which enables the capture of both local and global contextual information. RNAGenesis is composed of 32 Transformer blocks, each containing a multi-head attention mechanism and a feed-forward network (FFN). Positional encoding is provided by RoPE^69^, effectively representing both relative and absolute positional cues. Each multi-head attention module has 20 attention heads. To enhance pre-training efficiency at this scale, we integrate IO-aware FlashAttention^70^, a fast and memory-efficient exact attention mechanism. In the FFN, we employ two linear layers coupled with the SwiGLU activation function^71^, which merges the benefits of the Swish activation function and gated linear units (GLU). Residual connections link the Transformer modules, and layer normalization^72^ is applied inside each residual block to stabilize training and ensure well-behaved gradients at initialization.

#### Decoder

Besides encoding RNA sequences for understanding tasks, RNAGenesis has the unique capability of generating *de novo* RNA sequences. A Querying Transformer (QFormer)^23^, denoted by QFormer_*ϕ*_(·), is used to bridge the encoder and the decoder. It takes *K* trainable query token embeddings (*q*^1^,…, *q*^*K*^) as input, conducts cross-attention with the embeddings from the Encoder, and summarizes the information into *K* latent vectors (*z* = *z*^1^, *z*^2^,…, *z*^*K*^). *K* is a fixed number that does not depend on the input length *L*. Such a Q-Former preserves the key information for decoding while helping reduce the dimensionality by limiting *K*.

For decoding, we employ a standard causal Transformer^29^, denoted by Dec_*ψ*_. We first use a linear projection layer to convert the *K* latent vectors into a set of soft prompt embeddings with dimensions that match the decoder’s input size. Conditioned on these soft prompts, the decoder then autoregressively produces RNA sequence. With the pretrainedd RNAGenesis encoder, the decoder including the Q-Former and the causal Transformer is trained to reconstruct the input RNA sequences with the following loss function:

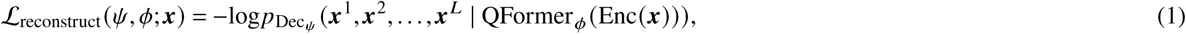

Taking the hybrid N-Gram tokenizer, the encoder backbone, and the decoder into consideration, the total model has 865.8M parameters.

#### Latent Diffusion

We leverage a continuous diffusion model^73^ to model the distribution of the latent variables *z*, which generates samples from the target distribution by a series of denoising processes. Specifically, we adopt a denoising network to predict the added noise *ϵ* from the noisy input 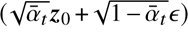, where *t* denotes the time step, 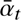 is the corresponding signal strength level, *z*_0_ is the uncorrupted latent vector, and *ϵ* is the added Gaussian noise. For the denoising score network Denoiser_*θ*_, we utilize transformer-based denoising networks following^74^. We adopt the *ϵ*-prediction scheme from Denoising Diffusion Probabilistic Models (DDPM)^73^ and employ the simplified training objective defined by

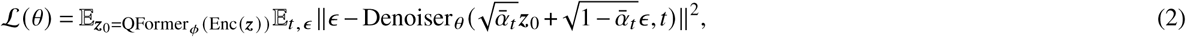

In our framework, the latent embeddings have fixed sizes, facilitating the training of diffusion models and allowing data to be treated as continuous, similar to images. Unlike other discrete diffusion models, our framework does not require truncating or padding sequences to a fixed length.

### Training and Finetuning

We pretrained RNAGenesis using a masked language modeling (MLM) objective. In this setup, unlabeled RNA sequences are corrupted and then reconstructed by the model. Each nucleotide in the RNA sequence is treated as a single token. We randomly selected 15% of the tokens in the training sequence. Among these, 80% are replaced with the special [MASK] token, 10% with a randomly chosen token from the vocabulary, and the remaining 10% are left unchanged. We used ordinary cross-entropy training loss function and our maximum sequence length is 1024. We use the Adam optimizer with *β*_1_ = 0.9, *β*_2_ = 0.999, and *ϵ* = 1 × 10^−8^. The initial learning rate is set to 1e-5, which increases to 1e-4 during the warmup phase and then gradually decays thereafter. We trained the RNAGenesis for 3 days on 16 NVIDIA Tesla A100-SXM4-80GB GPUs.

After pre-trained, RNAGenesis’s output embeddings can serve as a powerful sequence representation that has embedded structural and evolutionary information. Such a representation can be used as an enriched input to structural, functional, and engineering downstream tasks. We employed RNAGenesis for validation on 13 datasets provided by BEACON^15^, achieving state-of-the-art results on 10 of them. The incorporation of the CNN module significantly enhanced the performance of RNAGenesis in structural prediction tasks.

Following BEACON^15^, RNAGenesis incorporates three pipelines for different types of downstream tasks in BEACON and RNATx-Bench. For sequence-level tasks, we do not use the CLS token for prediction; instead, we apply an average readout function to the representations. Then representations are processed through an MLP layer to derive the sequence-level predictions. For tasks that demand resolution at the nucleotide level, each nucleotide’s representation is individually processed through an MLP to generate predictions specific to that nucleotide. To examine the relationships between nucleotides, a self-outer product of the nucleotide representations is calculated, resulting in a matrix that encapsulates pairwise interactions among nucleotides. This interaction matrix is subsequently processed through a simple ResNet to derive the final output. Except for the SSP task, which employed full fine-tuning, all other tasks utilized LoRA fine-tuning. For each task, we appropriately adjusted hyperparameters such as batch size, learning rate, and the LoRA ratio to optimize performance.

For sequence design, we followed RNAdiffusion^22^ and performed a comparative evaluation with RNA-FM. Specifically, we froze the pretrained encoder from RNAGenesis and trained the Q-former, modified from the ESM-2^75^ model architecture, along with the decoder, which uses the ProGen2-small^76^ architecture. The Q-former and decoder were trained from scratch on the ncRNA subset of the Ensembl database, comprising 1.1 million samples, over 2 epochs using two NVIDIA Tesla A100-SXM4-80GB GPUs. This training step encodes RNA sequences of varying lengths into fixed-size embeddings, with *K* = 16 and hidden dimension *D* = 160. Subsequently, a continuous diffusion model was trained using the same dataset for 2 epochs to model the distribution of the latent variables. After pretraining on the ncRNA dataset, the model was fine-tuned on the SELEX dataset A, using the same data-split strategy as AptaDiff^54^. The fine-tuning process was performed over 10 epochs. An identical pipeline was applied to RNA-FM to enable direct comparison.

### Inference-time Alignment

In this paper, we introduce two complementary strategies to enhance RNAGenesis for generating RNA sequences with desired properties, such as cleavage capability, secondary-structure similarity, and minimum free energy, which can be synergistically combined for optimal performance. The **Gradient Guidance** approach trains a reward model *R*_*ξ*_ on a labeled dataset {(**x**, *r*)} to predict rewards in the latent space via a Q-Former embedding **z** = *Q*_*ϕ*_(enc(**x**)), steering the denoising process by adjusting Denoiser_*θ*_’s noise prediction with a gradient term 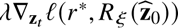. This method offers continuous, fine-grained control over the diffusion trajectory but may converge to local reward optima. In contrast, the **Beam Search** method, outlined in Algorithm 1, iteratively denoises ***B*** initial noisy latents over *T* timesteps using Denoiser_*θ*_ and 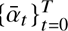, expanding each into *K* candidates and selecting the top ***B*** based on reward *R*. This discrete search excels at exploring diverse high-reward regions but lacks the gradient-based precision of guidance. By integrating Gradient Guidance’s smooth optimization with Beam Search’s broad exploration, RNAGenesis can leverage their strengths—precision and diversity—simultaneously, enhancing the efficiency and quality of reward-driven latent space sampling.

#### Gradient Guidance

To generate RNA sequences with desired properties, we require a reward model to predict the properties of new sequences. Suppose we have a labeled dataset {(**x**, *r*)}, where *r* represents the measured reward of interest, such as cleavage capability. We train a reward model (i.e., a neural network) *R*_*ξ*_ on the latent space to predict *r* given an input sequence **x.** The training objective is defined as

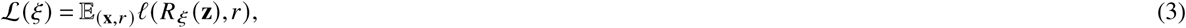

where *l* is a loss function, **z** = *Q*_*ϕ*_ (enc (**x**)) is the embedding obtained from the Q-Former, and *ξ* denotes the parameters of the reward model.

Subsequently, we employ the reward model to compute gradient-based guidance and integrate it into the denoising process of RNAGenesis. This guidance steers the random paths of the backward process toward regions of higher reward in a plug-and-play manner. Specifically, at step *t* of the backward diffusion process, given the current noisy sample **z**_*t*_, we estimate the clean sample as

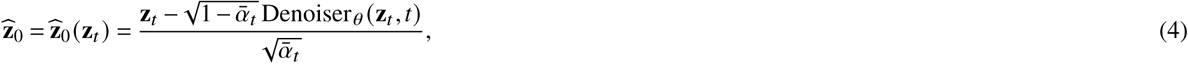

where Denoiser_*θ*_ (**z***_t_*, *t*) is the noise prediction model parameterized by *θ*. We then use 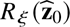 as a surrogate reward for the noisy sample **z**_*t*_. The gradient 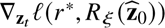 is incorporated into the predicted noise as follows, where *r*^∗^ is the target reward and *λ* is the guidance strength following previous works^77^:

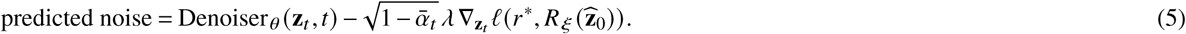

#### Beam Search

The pseudo algorithm of beam search for latent diffusion is summarized in Algorithm. 1 following previous works^78, 79^. Such a method leverages a latent diffusion model to generate high-reward samples by combining the generative power of diffusion processes with a guided beam search strategy. The algorithm operates by iteratively denoising an initial set of ***B*** noisy latent vectors 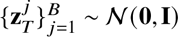 over *T* timesteps, guided by a noise scheduling parameter 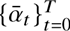. At each timestep *t*, the algorithm employs a trained denoiser Denoiser_*θ*_ to estimate the clean latent 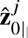, which is then used to sample the next latent 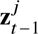. For *t* > 1, the process expands each beam into *K* candidate latents by adding controlled noise proportional to 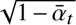, evaluates their estimated clean samples 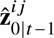 using a reward function *R*, and selects the top ***B*** candidates with the highest rewards to proceed to the next timestep. Note that the reward model could be non-differentiable, such as calculated secondary structure similarity and minimum free energy. This iterative refinement and selection process continues until *t* = 1, yielding a set of ***B*** final latents 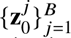, from which the highest-reward sample is returned. By integrating diffusion-based sampling with beam search optimization, the algorithm efficiently explores the latent space to produce samples that maximize the specified reward. In RNAGenesis with beam search strategy, the default setting of ***B*** and *K* are 1 and 8 respectively.

### Adapting RNAGenesis for Structural Tasks

RNAGenesis is adapted to structural tasks, including inverse folding, sequence-to-structure prediction, and *de novo* structure generation by combining it with structural modules and fine-tuning with LoRA^27^ and adapters^80^. The rank of LoRA size is set to 32 in the default setting. Generally, the sequence embeddings from RNAGenesis are fused with representations from other modules (e.g., GVP structure encoder) using a “project and concatenate” combiner module with simple MLPs following FoldFlow2^26^. If 2D features are needed for downstream tasks (e.g., IPA Transformer for structure prediction), the RNAGenesis embeddings undergo outer product to get the 2D features for fusion. We use LayerNorm^72^ for the fused embedding as it is essential to accommodate differently-scaled inputs.

#### GVP Encoder for RNA Structure Representation

To encode RNA tertiary structures, we adopt the Geometric Vector Perceptron (GVP) module from the RhoDesign architecture. The GVP encoder captures both vector- and scalar-valued features from the RNA backbone geometry. Vector features are computed from directional vectors between specific backbone atoms—most notably, vectors such as 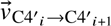, which describe the spatial orientation of nucleotide *i* relative to its neighbor. Each backbone atom (C4^′^, C1^′^, N1) is thus represented by a set of learned vector-valued embeddings reflecting local structural orientation. In parallel, scalar features are derived from dihedral angles involving seven backbone atoms (C4^′^, C1^′^, N1, C2, C5^′^, O5^′^, P), encoding local geometric configurations that are invariant to rigid-body transformations. These scalar and vector features are processed through GVP layers using linear projections, concatenation, and non-linearities based on the L2 norm. The resulting representations preserve rotation and reflection equivariance (for vector features) and invariance (for scalar features), and are passed to a downstream Transformer for sequence generation.

#### Transformer Decoder for Sequence Generation

The sequence decoder in RNAGenesis is part of a standard Transformer architecture composed of both an encoder and a decoder. The encoder receives inputs that include the structural features and a contact map (optional) derived from experimentally determined or RNAGenesis-predicted RNA structures. These contact maps, extracted using DSSR^81^, provide information on the spatial proximity of base pairs, allowing the model to incorporate long-range structural dependencies. The encoder applies multi-head attention mechanisms to integrate this spatial information, producing a structurally enriched representation of the RNA backbone. The decoder then autoregressively generates nucleotide sequences by attending to the encoder’s output and previously generated tokens, enabling context-aware sequence generation conditioned on 3D structure. The Transformer architecture used in RNAGenesis includes three encoder and three decoder layers, each with four attention heads and an embedding dimension of 512. Attention dropout is set to 0.1. The model is trained using a cross-entropy loss function, and optimized using data derived from both experimentally resolved RNA structures (RNASolo^38^). During training, contact maps serve as auxiliary inputs to enhance structural inductive bias, while the decoder learns to generate nucleotide sequences that are compatible with the given tertiary backbone geometry.

**Algorithm 1.**
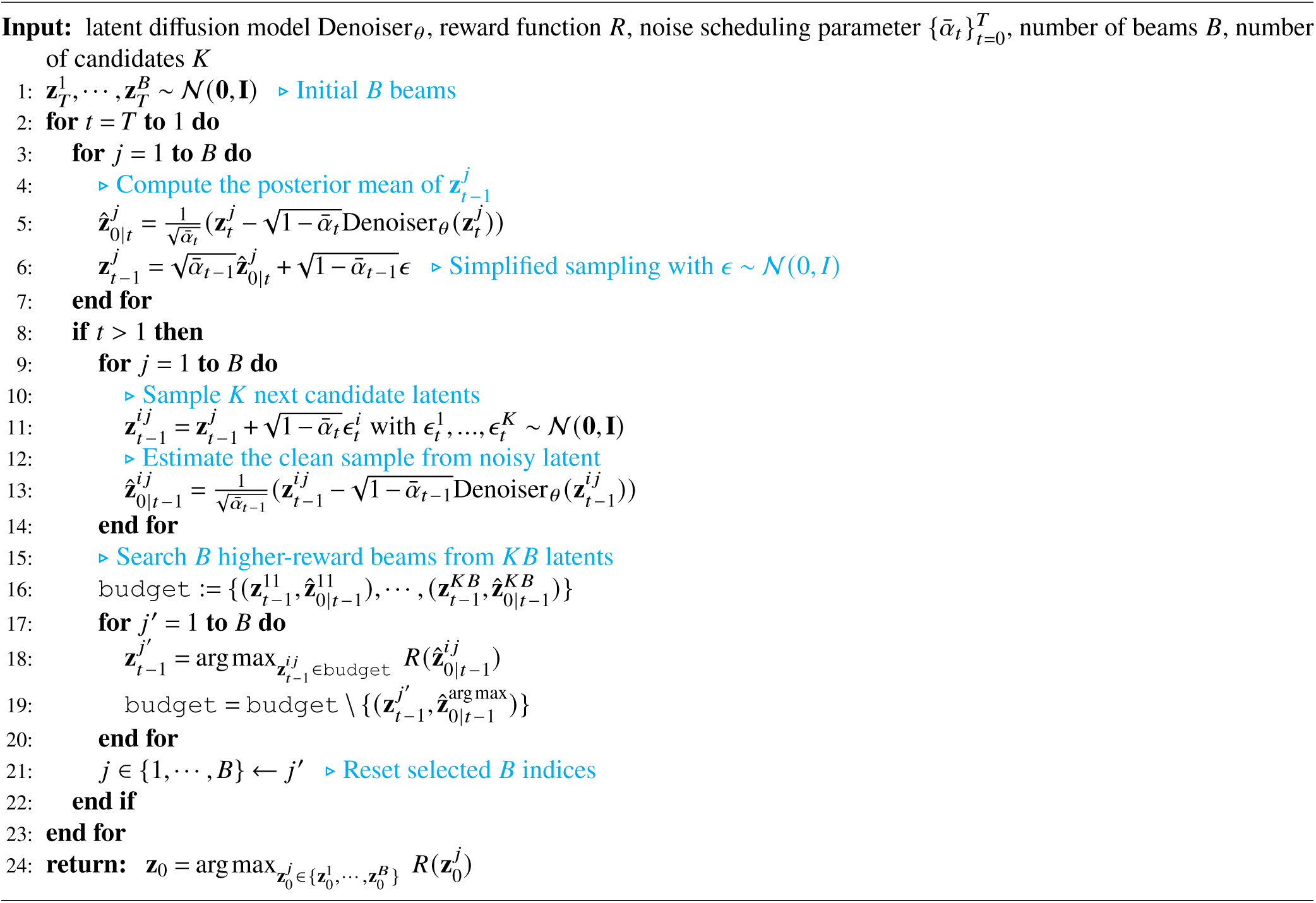
Diffusion Latent Beam Search with 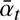

#### Structure Decoder via IPA Transformer and SE(3) Flow Matching

For RNA 3D structure generation, RNAGenesis adopts a structure decoder based on the *Invariant Point Attention (IPA) Transformer*^82^, originally introduced in AlphaFold. The IPA module is specifically designed to capture spatial relationships between residues (or nucleotides) in a way that is equivariant to 3D rotations and translations. In RNAGenesis, the IPA Transformer operates over RNA backbone *frames*, where each nucleotide is represented by a rigid body frame defined by the *C*3^′^, *C*4^′^, and *O*4^′^ atoms. This compact representation enables the model to reason over the spatial configuration of the RNA chain without modeling all 13 backbone atoms explicitly.

To generate full backbone structures, RNAGenesis follows the SE(3) flow matching framework, as in RNA-FrameFlow^24^. Each nucleotide frame is parameterized by a 3D translation vector (placing *C*4^′^) and a 3×3 rotation matrix (orienting the *C*3^′^-*C*4^′^-*O*4^′^ triangle). Rather than predicting each atom’s coordinates independently, the model learns a vector field that maps noisy initial frames—sampled from ***N***(0,**I**) × ***u***(*SO*(3)) —toward the true distribution of RNA structures. This interpolation is performed via geodesic paths on the *SE*(3) manifold using exponential and logarithmic maps. The IPA Transformer provides attention-based updates conditioned on structural context, guiding the denoising process to ensure local coherence and global consistency. After frame generation, the full-atom RNA structure is reconstructed by autoregressively placing the remaining atoms using predicted torsion angles, completing the all-atom model.

### Baselines

Table S2 provides an overview of related RNA foundation models, predominantly based on encoder-only transformer architectures and pre-trained using masked language modeling (MLM) objectives, often with 15–25% masking ratios, alongside other tasks. These models vary significantly in size and pre-training data: for instance, RNABERT^5^ and UTR-LM^7^ are smaller (<10M parameters) with focused datasets like RNAcentral and 5’ UTRs, while larger models like Uni-RNA^11^ and AIDO.RNA^11^ scale up to 400M and 1.6B parameters, respectively, leveraging extensive datasets such as RNAcentral and nt & Genome Warehouse (1B sequences) or RNAcentral v24.0 (42M ncRNA sequences). Most models, including RNA-FM^6^, SpliceBERT^8^, and RiNALMo^10^, use base-level tokenization and MLM, with some like CodonBERT^14^ and CaLM^83^ adopting codon-based tokenization for specific applications. In contrast, RNAGenesis (ours), with an encoder-decoder transformer architecture (∼ 1B parameters), is pre-trained on RNAcentral v24.0 using a hybrid N-Gram MLM (30% masking), enabling robust generative capabilities. Unlike the encoder-only baselines focused on representation learning, RNAGenesis excels in RNA sequence generation, further enhanced by integrating gradient guidance and beam search during inference. Gradient guidance steers the denoising process toward high-reward regions using a reward model *R*_*ξ*_, while beam search explores diverse high-reward latents, combining continuous optimization with discrete search to efficiently generate RNA sequences with desired properties like cleavage capability.

For structural tasks, we consider the RhoDesign Ribologic, Learna, gRNAde, Rdesign, and eM2dRNAs for inverse folding; RNA-frameflow for *de novo* structure generation; RhoFold+, FARFAR2, DRFold, and AlphaFold2 for structure prediction from sequence. We use the provided model checkpoints in their respective repositories, that is https://github.com/ml4bio/RhoDesign for RhoDesign, https://github.com/automl/learna for LEARNA, https://github.com/wuami/RiboLogic for RiboLogic; https://github.com/chaitjo/geometric-rna-design for gR-NAde, https://github.com/A4Bio/RDesign for RDesign, https://github.com/iARN-unex/eM2dRNAs for eM2dRNAs, https://github.com/ml4bio/RhoFold for RhoFold+, https://rosie.rosettacommons.org/farfar2 for FARFAR2, https://github.com/leeyang/DRfold for DRFold, https://github.com/google-deepmind/alphafold3 for AlphaFold3, and https://github.com/rish-16/rna-backbone-design for RNA-FrameFlow. Default parameters for each model were used without any modifications, and no additional tuning or parameter adjustments were performed.

For RNA therapeutic tasks and Human UTR variants effect prediction tasks, RNAGenesis is compared with representative open-source RNA/DNA models, including RNA-FM ^6^, Evo^84^, and Evo2^16^. For these prediction tasks, we mounted a classification head, consisting of three self-attention layers followed by a two-layer MLP, onto the sequence embeddings generated by each model.

### Surface Plasmon Resonance Assay

Surface plasmon resonance (SPR) experiments were conducted using the Biacore X100 instrument (Cytiva). The target protein for dataset A was insulin-like growth factor-binding protein 3 (IGFBP3). Aptamers previously identified through SELEX wet-lab screening were used as positive controls. All tested sequences are provided in the Supplementary Information. Aptamers were synthesized according to their sequence information and obtained as solid powders (GenScript, Nanjing, China).

SPR experiments were performed using a Biacore sensor chip CM5 (29149604, Cytiva). IGFBP3 protein was immobilized onto the chip surface following the manufacturer’s standard amine coupling protocol. The running buffer was 1× HBS-EP+ (pH 7.4; BR100669, Cytiva) supplemented with 5 mM MgCl_2_. Aptamer candidates were diluted in phosphate-buffered saline (PBS) containing 5 mM MgCl_2_ to specific concentrations and injected over the chip surface at a flow rate of 30 *μ*L/min. Each binding cycle consisted of 120 s association and 120 s dissociation phases.

To regenerate the sensor surface between cycles, 1.5 M NaCl supplemented with 5 mM NaOH was used as the regeneration buffer (flow rate: 30 *μ*L/min; duration: 60 s). Binding kinetics, including the equilibrium dissociation constant (*K*_*D*_), association rate constant (*k*_*a*_ or *k*_on_), and dissociation rate constant (*k*_*d*_ or *k*_off_), were calculated using the Biacore Evaluation Software (version 2.0.1.201). Binding curves were post-processed and smoothed using RStudio.

### Guide RNA Variant Vector Construction

Gene fragments containing *sp*Cas9 spacer sequences targeting *B2M* (5’-CGCGAGCACAGCTAAGGCCA-3’) and *EGFP* (5’-TAGGTGGCATCGCCCTCGCC-3’), along with guide RNA variants, were synthesized by Twist Biosciences. These fragments were amplified using Phanta Flash Master Mix (New England Biolabs, NEB). Amplified products were gel-purified using the Monarch DNA Gel Extraction Kit (NEB) and subsequently inserted into a *BsmBI* and *EcoRI* digested lentiCRISPR v2 vector backbone (Addgene, cat. no. 52961) via Gibson Assembly with the NEBuilder HiFi DNA Assembly Master Mix (NEB). The ligated plasmid products were transformed into Stbl3 Chemically Competent *E. coli* (Zymo Research). All constructs were validated by Sanger sequencing (Azanta).

### Mammalian Cell Culture

HEK 293T cells (ATCC) were cultured in Dulbecco’s Modified Eagle’s Medium (DMEM; Gibco, cat. no. 11965-092) supplemented with 10% (v/v) fetal bovine serum (Gemini Bio, cat. no. 100-106-500) and 1% penicillin-streptomycin (Gibco, cat. no. 15-140-122). Cells were maintained in a humidified incubator at 37°C with 5% CO_2_.

### CRISPR-Cas9 Cleavage Assays

#### EGFP Plasmid Reporter Cleavage Assay

HEK293T cells were seeded into 96-well plates (Corning) at a density of 2.5 × 10^4^ cells per well, 24 hours prior to transfection. For each well, a mix containing 10 ng of a pcDNA3.1-EGFP plasmid, 10 ng of a pcDNA3.1-mCherry plasmid, and 90 ng of the modified lentiCRISPR v2 plasmid expressing a guide RNA variant was transfected using PEI MAX (Polysciences, cat. no. 24765). At 48 hours post-transfection, EGFP and mCherry fluorescence levels were measured by flow cytometry using a CytoFLEX analyzer (Beckman Coulter).

#### Endogenous B2M Cleavage Assay

HEK293T cells were seeded into 96-well plates (Corning) at a density of 1.5 × 10^4^ cells per well, 24 hours prior to transfection. For each well, 90 ng of the modified lentiCRISPR v2 plasmids with protospacer 5’-CGCGAGCACAGCTAAGGCCA-3 and scaffold variants were individually co-transfected with 5 ng of a pcDNA3.1-GFP plasmid (as a transfection marker) using PEI MAX. At 72 hours post-transfection, cells were harvested and stained with an anti-human B2M-APC antibody (BioLegend, cat. no. 316311) in PBS containing 5% BSA. Surface B2M expression and EGFP expression were quantified by flow cytometry on a CytoFLEX analyzer (Beckman Coulter).

#### Endogenous AAVS1 Editing Assay

To compare sgRNA-scaffold variants at the genomic AAVS1 locus, the protospacer 5’-GGGGCCACTAGGGACAGGAT-3’ was cloned upstream of each scaffold into lentiCRISPR v2 (Addgene #98293). HEK293T cells were plated as above and transfected with 50, 100 or 250 ng of the respective plasmid. Medium was refreshed 12 h later, and cells were collected 72 h post-transfection. Genomic DNA was extracted with QuickExtract DNA Extraction Solution (Lucigen, QE09050), the target region was PCR-amplified, and analysed with TIDE (https://tide.nki.nl/) to quantify indel frequencies.

### Prime Editing PEAR Reporter Assay

To assess prime editing efficiency, we utilized the previously developed PEAR reporter system. HEK293T cells were seeded in 96-well plates at a density of 1.5 × 10^4^ cells per well, 24 hours prior to transfection. Each well was co-transfected with 30 ng of the PEAR-GFP-2in1 plasmid (Addgene, cat. no. 177186) containing the target site for a specific pegRNA variant, along with 70 ng of the PEMAX plasmid (Addgene, cat. no. 191102), which expresses the prime editor enzyme. Transfections were performed using PEI MAX. At 72 hours post-transfection, the percentage of EGFP-positive cells, indicating successful prime editing, was measured relative to mScarlet expression by flow cytometry on a CytoFLEX analyzer (Beckman Coulter).

### Base Editing Reporter Assay

The BEAR (Base Editing Activity Reporter) system was used to evaluate the editing efficiency of different gRNA scaffold variants, as described in previous works^85^. HEK293T cells were seeded into 48-well plates at 1.5 × 10^4^ cells per well 24 h before transfection. Cells were co-transfected with 60 ng of a single-plasmid BEAR construct (encoding both the sgRNA and the GFP reporter) and either 60 ng or 120 ng of a cytosine or adenine base editor (Addgene, #163008, #174127) expression plasmid, using Lipofectamine 3000 (Thermo Fisher Scientific) following the manufacturer’s protocol. Culture medium was replaced 12 h post-transfection. Seventy-two hours after transfection, cells were harvested and analyzed on a flow cytometer; the gating strategy is shown in Supplementary Fig. S14.

## Data Availability

All data used in this study are publicly available, with details of their usage provided in the Methods section. The RNAGenesis dataset is sourced from RNACentral (https://rnacentral.org/). Fine-tuning datasets for the Beacon benchmark are curated at https://github.com/terry-r123/RNABenchmark. Datasets for structural tasks are obtained from RNASolo (https://rnasolo.cs.put.poznan.pl/), CASP (https://predictioncenter.org/index.cgi), and RNA Puzzles (https://www.rnapuzzles.org/). Fine-tuning datasets for ASO, CircRNA, siRNA, and shRNA are from supplementary data of previous papers^45–48^. Fine-tuning datasets for human UTR effect predictions are from https://doi.org/10.1038/s41467-024-46795-7, https://doi.org/10.1186/s13073-023-01277-1, and https://doi.org/10.1093/bib/bbae248. The training dataset for aptamer design is available at https://github.com/wz-create/AptaDiff/blob/master/data/raw_data/datasetA_IGFBP3_P6.csv. The training dataset for sgRNA scaffold design is accessible via https://www.cell.com/cell-chemical-biology/fulltext/S2451-9456(23)00184-8#mmc3. Additional datasets are available from the Protein Data Bank (https://www.rcsb.org/).

## Code availability

The source code of this study is freely available at GitHub (https://github.com/zaixizhang/RNAGenesis) to allow for replication of the results of this study. This study’s checkpoints and configs are available at Zenodo (https://zenodo.org/records/15192633). The project website is at https://zaixizhang.github.io/ZaixiZhang_files/RNAGenesis.html.

## Acknowledgements

None

## Author contributions statement

Z.X.Z, R.F.J., L.L.C, L.C., and M.D.W conceived the study and designed the experiments. L.L.C, Y.K.Z., and G.W.Z. performed the dataset analyses and developed the pre-trained models. Z.X.Z., L.L.C., R.F.J., Y.K.Z., G.W.Z.,Y.Q.G, Y.K.Y., and K.X.H. conducted the downstream computational experiments and collected the data. G.X.X., D.Y., Y.Q.F., X.T.W., and J.X.Z. conducted the wet-lab experiments and collected the data. Z.X.Z, R.F.J., Y.J.Y., Q.R.Y., and L.C. analyzed the data and interpreted the results. Z.Y.X., W.N.E., R.H.Z., X.M.Z., L.C., and M.D.W. supervised the project and provided critical intellectual input. Z.X.Z, L.L.C., and R.F.J. wrote the initial draft of the manuscript. W.N.E., X.M.Z., L.C., and M.D.W. revised the manuscript. All authors reviewed and approved the final version of the manuscript.

## Competing interests

The authors declare no competing interests.

## Additional information

**Correspondence and requests for materials** should be addressed to Ruhong Zhou, Xiaoming Zhang, and Mengdi Wang, Le Cong.

## Materials and Methods

### Pretrain Data Distributions

We leverage the preprocessed dataset from RNACentral^12^ for the pretraining, which contains a total of 42 million sequences. To reduce the impact of data redundancy on the model, we use MMSeq2 to cluster the sequences with a similarity threshold of 90%. After clustering, the total number of cluster centers is reduced to 6.4 million. Figure S1 illustrates the distribution changes of RNA samples based on type before and after clustering. In total, 22 RNA types are represented. The most significant change is observed in rRNA, which initially accounted for 70.78% of the dataset before clustering but is reduced to 21.04% after clustering. This suggests that rRNA sequences have a high level of redundancy, and clustering effectively reduce their over representation, improving dataset balance. In contrast, lncRNA, which makes up 7.79% before clustering, increases to 36.03% after clustering, indicating that lncRNA sequences are more diverse and retain a larger proportion of unique representatives. Other RNA types, such as tRNA, misc RNA, and sRNA, also show an increase in their proportion after clustering, reflecting the removal of redundant sequences from dominant categories like rRNA. Smaller RNA types, including guide RNA, RNase MRP RNA, and Y RNA, exhibit minimal changes, suggesting relatively low redundancy in those categories. This transformation enhances the dataset’s quality by reducing redundancy while preserving sequence diversity, ensuring a more representative and balanced distribution of RNA types. During model pretraining, we first randomly select cluster centers and then sample from the corresponding clusters to form a batch.

### Descriptions of the BEACON Benchmark Tasks

#### Structure Prediction

##### Secondary structure prediction (SSP)

This task utilizes the bpRNA-1m database^86^, which provides comprehensive annotations for over 100,000 single-molecule RNA structures, as the primary source of data for SSP. It focuses on identifying base-paired regions (stems) as well as unpaired regions such as loops, bulges, and junctions within RNA sequences. The ground-truth labels are represented by a matrix, where each element indicates whether a base pair is formed between nucleotide positions.

##### Contact Map Prediction (CMP)

This task utilizes a dataset derived from non-redundant RNA 3D structures, as compiled by^87^ and adopted in^88^. It involves identifying nucleotide pairs that are spatially adjacent in the tertiary structure of RNA molecules. Each pair is annotated with a binary label, where 1 denotes a contact—defined as a distance of less than 8 °A between nucleotides—and 0 otherwise.

##### Distance Map Prediction (DMP)

The dataset used for this task is the same as that employed in the CMP task, derived from a curated collection of non-redundant RNA 3D structures. The objective of DMP is to estimate the spatial distances between all nucleotide pairs in an RNA molecule. The target is represented as a continuous matrix, where each element encodes the physical distance between two nucleotides in the sequence.

##### Structural Score Imputation (SSI)

This task utilizes a dataset introduced in^89^, which is derived from icSHAPE profiling experiments conducted on the HEK293 human cell line. The goal of SSI is to recover missing structural signals by predicting nucleotide-level structural scores. Each nucleotide is assigned a continuous value, representing an experimentally determined structural property. To simulate data incompleteness, 30% of the scores are randomly masked during training, and the test set includes 3,095 RNA fragments with downsampled missing values.

#### RNA Function Prediction

##### Splice Site Prediction (SPL)

This task builds upon the dataset introduced by Jaganathan et al.^90^, for splice site classification. The SPL task involves identifying the functional role of each nucleotide in a given sequence. Specifically, each position is categorized as an acceptor, donor, or neither.

##### APA Isoform Prediction (APA)

This task makes use of a dataset curated by Bogard et al.^91^, which comprises over 3 million APA reporter gene sequences. From this collection, 228,000 sequences were selected for training and evaluation. The APA task aims to estimate the relative usage of the proximal polyadenylation site (PAS) within the 3’ untranslated region (3’ UTR) of RNA variants. The target for each sequence is a continuous value, representing the proportion of isoforms utilizing the proximal PAS.

##### Non-coding RNA Function Classification (ncRNA)

The dataset used for this classification task is compiled from multiple well-established sources, including GENCODE, circBase, and Rfam, as integrated and described in^92, 93^. It encompasses a wide spectrum of non-coding RNA types with diverse biological functions. The objective of ncRNA is to categorize each ncRNA molecule into its corresponding functional class, such as microRNAs (miRNAs), long non-coding RNAs (lncRNAs), or small interfering RNAs (siRNAs). Each RNA sequence is annotated with a categorical label, denoting its functional identity.

##### Modification Prediction(Modif)

This task leverages a dataset compiling 20 epi-transcriptome profiles covering 12 distinct RNA modification types. The data is derived from 15 high-resolution sequencing technologies and includes over 300,000 annotated modification sites. The objective of the Modif task is to identify the specific type of RNA chemical modification present at each site within a given RNA sequence. Each site is assigned a categorical label, corresponding to one of the twelve most frequent RNA modification types.

##### Mean Ribosome Loading (MRL)

This task utilizes the dataset published by Reid et al.^94^, which contains 91,519 annotated 5’ UTR sequences along with their engineered variants. The goal of the MRL task is to quantify the level of mRNA translation efficiency by predicting the MRL value for each input sequence. The target variable is a continuous value, reflecting the mean number of ribosomes loaded onto an mRNA molecule.

#### Engineering Prediction

##### Vaccine Degradation Prediction (VDP)

This task makes use of the dataset released through the “Stanford OpenVaccine” competition on Kaggle, supplemented with experimental measurements from the RNA design platform Eterna^95^. The dataset includes 6,043 synthetic RNA constructs with annotated degradation profiles under various environmental conditions. The objective of the VDP task is to estimate the stability and shelf life of RNA-based vaccines by modeling their degradation characteristics. For each nucleotide in a sequence, the target is a vector, representing three molecular properties associated with degradation dynamics. Model accuracy is evaluated using the Mean Columnwise Root Mean Squared Error (MCRMSE) metric. **Programmable RNA Switches (PRS):** This task is based on a dataset curated and analyzed by Angenent-Mari et al.^96, 97^, comprising 91,534 experimentally validated toehold switches tested in vivo. These synthetic RNA constructs target 23 viral genomes and 906 human transcription factors, with corresponding GFP fluorescence measurements quantifying their regulatory activity. The PRS task aims to identify RNA sequences capable of dynamic structural and functional shifts in response to molecular signals. For each sequence, the target is a continuous vector, capturing ON, OFF, and ON/OFF state expression levels.

##### CRISPR On-Target Prediction(CRI-On)

The dataset is provided by Chuai et al.^98^, containing approximately 15,000 single-guide RNAs (sgRNAs) targeting 1,071 genes across four distinct human cell lines. The CRI-On task focuses on estimating the gene knockout efficiency of sgRNAs guided by Cas proteins at specific genomic loci. Each sgRNA is assigned a continuous value, reflecting its measured editing efficacy.

##### CRISPR Off-Target Prediction (CRI-Off)

This task is based on the off-target evaluation dataset compiled by Chuai et al.^98^, which includes approximately 160,000 potential off-target genomic loci associated with 30 distinct sgRNAs across multiple human cell types. The CRI-Off task aims to estimate the frequency of unintended mutations induced by CRISPR at non-targeted genomic sites. Each site is assigned a continuous target value, representing the observed off-target editing frequency.

### Descriptions of RNATx-Bench Tasks

#### Antisense Oligonucleotide Prediction (ASO)

This task evaluates the ability of models to predict the inhibitory efficacy of antisense oligonucleotides (ASOs) targeting specific disease-related genes. The dataset consists of 34,773 experimentally validated ASO sequences paired with inhibition rates across four clinically relevant targets: SOD1, K-RAS, APOL1, and SNHG14. **SOD1** is a validated target in amyotrophic lateral sclerosis (ALS), with the ASO therapy tofersen (Qalsody) approved by the FDA in 2023 for reducing SOD1 protein and neurofilament light chain levels—an indicator of neuronal injury^99^. **K-RAS**, a key oncogene in lung, colorectal, and pancreatic cancers, was long considered “undruggable,” but ASOs like AZD4785 have recently shown promise in inhibiting its mRNA to suppress tumor growth^100^. **APOL1** variants increase kidney disease risk, especially in individuals of African descent; ASOs targeting APOL1 reduce proteinuria and preserve kidney function in preclinical studies^101^. **SNHG14** regulates UBE3A expression, disrupted in Angelman syndrome; ASOs against SNHG14 can reactivate paternal UBE3A in neurons, offering a therapeutic strategy^102^. The goal is to regress the inhibition score based on sequence features. Ground truth labels are continuous values representing the experimentally measured inhibition rate for each ASO.

#### Circular RNA Regulatory Interaction (CircRNA)

This task involves predicting the functional interactions of circular RNAs (circRNAs), including their binding with RNA-binding proteins (RBPs) and microRNAs (miRNAs). The dataset contains over 4,000 experimentally validated interactions. The goal is to classify whether a given circRNA binds to a particular RBP or miRNA. Ground truth labels are binary: 1 for a confirmed interaction and 0 otherwise.

#### RNA Aptamer Design and Evaluation (Aptamer)

This generative task assesses the capacity of models to design functional RNA aptamers that match experimentally derived SELEX sequences. Models are evaluated on four criteria: sequence similarity (edit distance), GC content, minimum free energy (MFE), and secondary structure similarity. A subset of top-performing generated sequences is also assessed using 3D structure prediction tools such as AlphaFold3 to evaluate structural confidence and folding plausibility.

#### Small Interfering RNA Efficacy Prediction (siRNA)

This task benchmarks models on predicting the silencing efficacy of siRNA sequences using two public datasets: Takayuki (N = 702) and Huesken (N = 2,361). The objective is to classify high-vs. low-efficacy siRNAs, with ground truth labels derived from experimental knockdown efficiency. Performance is evaluated using AUROC and AUPRC metrics.

#### Short Hairpin RNA Efficacy Prediction (shRNA)

Similar to siRNA, this task involves predicting the efficacy of shRNA sequences in gene knockdown experiments. The dataset includes 2,076 labeled examples with experimentally measured silencing efficiency. Labels are continuous scores, and models are evaluated using correlation coefficients (Pearson and Spearman).

#### UTR Variant Pathogenicity Classification (UTR-P)

This task involves classifying human 5’ and 3’ UTR variants as either pathogenic or benign. It uses variant annotations from inherited retinal disease (IRD) and ClinVar datasets (N = 1,547 and 6,265, respectively). Ground truth labels are binary pathogenicity assignments, and performance is measured using AUROC and AUPRC.

#### UTR Variant Regulatory Effect Classification (UTR-Reg)

This task leverages data from Massively Parallel Reporter Assays (MPRA) to classify UTR variants as up-regulating, down-regulating, or neutral based on their effect on gene expression. The dataset comprises 34,404 variants from MPRAs in HEK293 and HeLa cells. Ground truth labels are derived from experimental log-fold change (lnfc) values and binned into three categories. Models are evaluated using AUROC and AUPRC per class.

### Fine-tuning Settings of the BEACON benchmark

#### Secondary Structure Prediction

Secondary Structure Prediction identifies paired regions and unpaired regions within RNA molecules. We construct pairwise representations by employing outer concatenation on the outputs of the language model, wherein the representation of nucleotide *j* is concatenated with that of nucleotide *i* for each nucleotide pair (*i*, *j*). In the prediction head, this concatenated representation is initially projected linearly into a 64-dimensional embedding space. Subsequently, it passes through two ResNet-2D blocks and a convolutional layer, all of which utilize 64 kernels of size 3. Each layer is accompanied by instance normalization and ReLU activation. To train the model, we use AdamW optimizer with weight decay 0.01. For the secondary structure prediction task, we conducted full fine-tuning, using a batch size of 16 and a maximum learning rate of 2.5 × 10^−4^. When comparing our method with others in the BEACON benchmark, we adopt the *F*_1−0.5_ metric. For comparisons with RiNALMO and AIDO.RNA, we employ the *F*_max_ metric and apply the same post-processing techniques as those used in these methods.

#### Contact Map Prediction

Contact Map Prediction is a three dimension structure prediction task. For this task, we utilize the same prediction head as in the secondary structure prediction task and evaluate the model using Top-L precision. The key difference between SSP task is that we employ LoRA fine-tuning with batch size of 8 and maximum learning rate of 3 × 10^−4^.

#### Distance Map Prediction

Distance Map Prediction estimates the physical distances between pairs of nucleotides within an RNA molecule. For this task, we utilize the same prediction head as in the secondary structure prediction task and evaluate the model using the *R*^2^ metric. We employ LoRA fine-tuning with batch size of 2 and maximum learning rate of 3 × 10^−4^.

#### Structural Score Imputation

Structural Score Imputation predicts missing structural information within RNA molecules, where each nucleotide is assigned an experimentally derived structural score. This task is a nucleotide-level regression problem, evaluated using the *R*^2^ metric. During the fine-tuning process, the structure information is first projected through a linear layer to match the dimensionality of the hidden layer representation. Then, the two representations are concatenated and mapped to a 640-dimensional space by the input layer of the prediction head. This is followed by a two-layer MLP with a hidden dimension of 256, accompanied by ReLU activation. The model is trained and evaluated exclusively on regions where structural information is missing. To train the model, we employ the AdamW optimizer with a weight decay of 0.01. To mitigate overfitting, we apply a dropout rate of 0.2 to the prediction head. LoRA fine-tuning is adopted, using a batch size of 4 and a maximum learning rate of 5 × 10^−5^.

#### Splice Site Prediction

Splice Site Prediction categorizes each nucleotide within a sequence into one of three classes: acceptor, donor, or neither. This task is a nucleotide-level three-class classification problem, evaluated using the Top-*k* accuracy metric. In the prediction head, the sequence representation is mapped to a 640-dimensional space by the input layer of the prediction head. This is followed by a two-layer MLP with a hidden dimension of 256, accompanied by ReLU activation. To train the model, we employ the AdamW optimizer with a weight decay of 0.01. To mitigate overfitting, we apply a dropout rate of 0.2 to the prediction head. LoRA fine-tuning is adopted, using a batch size of 128 and a maximum learning rate of 1 × 10^−5^.

#### APA Isoform Prediction

APA Isoform Prediction predicts the usage ratio of the proximal polyadenylation site (PAS) in the 3’ untranslated region (3’ UTR) for each variant, recorded in target. This task is a sequence-level regression problem, evaluated using the *R*^2^ metric. Notably, instead of using the representation of the CLS token for prediction, we utilize the mean representation of all tokens in the sequence. In the prediction head, the sequence representation passes through a two-layer MLP with a hidden dimension of 128, accompanied by ReLU activation. To train the model, we employ the AdamW optimizer with a weight decay of 0.01. To mitigate overfitting, we apply a dropout rate of 0.1 to the prediction head. LoRA fine-tuning is adopted, using a batch size of 64 and a maximum learning rate of 1 × 10^−4^.

#### Non-coding RNA Function Classification

Non-coding RNA Function Classification classifies ncRNA molecules into categories like miRNAs, lncRNAs, and siRNAs. This task is a sequence-level thirteen-class classification problem, evaluated using the Accuracy metric. Notably, instead of using the representation of the CLS token for prediction, we utilize the mean representation of all tokens in the sequence. In the prediction head, the sequence representation passes through a two-layer MLP with a hidden dimension of 128, accompanied by ReLU activation. To train the model, we employ the AdamW optimizer with a weight decay of 0.01. To mitigate overfitting, we apply a dropout rate of 0.1 to the prediction head. LoRA fine-tuning is adopted, using a batch size of 16 and a maximum learning rate of 1 × 10^−4^.

#### Modification Prediction

Modification Prediction predicts twelve widely occurring types of RNA modifications from a given RNA sequence. This task is a sequence-level multi-label binary classification problem with twelve labels, evaluated using the AUC metric. In the prediction head, the sequence representation is transformed through a linear layer to match the number of labels. LoRA fine-tuning is adopted, using a batch size of 64 and a maximum learning rate of 1 × 10^−4^.

#### Mean Ribosome Loading

Mean Ribosome Loading estimates the MRL value for a given sequence, where the target represents the degree of mRNA translation activity into proteins. This task is a sequence-level regression problem, evaluated using the *R*^2^ metric. Notably, instead of using the representation of the CLS token for prediction, we utilize the mean representation of all tokens in the sequence. In the prediction head, the sequence representation passes through a two-layer MLP with a hidden dimension of 128, accompanied by ReLU activation. To train the model, we employ the AdamW optimizer with a weight decay of 0.01. To mitigate overfitting, we apply a dropout rate of 0.1 to the prediction head. Full fine-tuning is adopted, using a batch size of 8 and a maximum learning rate of 3 × 10^−5^.

#### Programmable RNA Switches

Programmable RNA Switches involves the identification of synthetic RNA molecules capable of altering their conformation and function in response to specific signals. This task is a sequence-level multi-label regression problem with three labels, evaluated using the *R*^2^ metric. In the prediction head, the sequence representation is transformed through a linear layer to match the number of labels. LoRA fine-tuning is adopted, using a batch size of 32 and a maximum learning rate of 1 × 10^−4^.

#### CRISPR On-Target Prediction

CRISPR On-Target Prediction evaluates the efficiency of single-guide RNAs (sgRNAs) directed by Cas proteins in gene editing at specific target sites. This task is a sequence-level regression problem, evaluated using the *Spearman* metric. Notably, instead of using the representation of the CLS token for prediction, we utilize the mean representation of all tokens in the sequence. In the prediction head, the sequence representation passes through a two-layer MLP with a hidden dimension of 128, accompanied by ReLU activation. To train the model, we employ the AdamW optimizer with a weight decay of 0.01. To mitigate overfitting, we apply a dropout rate of 0.1 to the prediction head. Lora fine-tuning is adopted, using a batch size of 32 and a maximum learning rate of 1 × 10^−4^.

#### CRISPR Off-Target Prediction

CRISPR Off-Target Prediction predicts the likelihood and frequency of CRISPR-induced mutations at unintended genomic locations. This task is a sequence-level regression problem, evaluated using the *Spearman* metric. Notably, instead of using the representation of the CLS token for prediction, we utilize the mean representation of all tokens in the sequence. During the fine-tuning process, “sgRNA” and “target” are both used as inputs to the model. Then, the two representations are concatenated and mapped to a 640-dimensional space by the input layer of the prediction head. This is followed by a two-layer MLP with a hidden dimension of 256, accompanied by ReLU activation. To train the model, we employ the AdamW optimizer with a weight decay of 0.01. To mitigate overfitting, we apply a dropout rate of 0.2 to the prediction head. LoRA fine-tuning is adopted, using a batch size of 32 and a maximum learning rate of 1 × 10^−4^.

### Fine-tuning Settings of RNATx-Bench

#### Antisense Oligonucleotide Prediction (ASO)

This task predicts the inhibition efficiency of antisense oligonucleotides targeting disease-associated genes. It is a sequence-level regression problem evaluated using Pearson and Spearman correlation coefficients. We extract the token-wise embeddings from the last hidden layer of RNAGenesis and compute the mean-pooled sequence representation. A two-layer MLP with a hidden dimension of 128 and ReLU activation is applied as the prediction head. The model is fine-tuned end-to-end using the AdamW optimizer with a learning rate of 1 × 10^−4^, weight decay of 0.01, and dropout of 0.1. We train for 100 epochs with a batch size of 32.

#### Circular RNA Interaction Prediction (circRNA)

This task involves binary classification of circRNA interactions with microRNAs or proteins. It is framed as a sequence-level classification problem and evaluated using AUROC and AUPRC. As with ASO, we mean-pool token embeddings from the last layer of RNAGenesis and pass them through a two-layer MLP classifier (hidden dimension 128, ReLU). Fine-tuning is performed using AdamW with a learning rate of 1 × 10^−4^, weight decay of 0.01, and dropout rate of 0.1. The model is trained with a batch size of 32 for 100 epochs.

#### shRNA Efficacy Prediction

The task is to predict gene silencing efficiency of short hairpin RNAs. It is treated as a regression problem with evaluation via Pearson and Spearman correlations. We adopt a similar mean-pooling plus MLP setup as in ASO, with a hidden dimension of 128 and ReLU activation. Fine-tuning is conducted with AdamW, learning rate 1 × 10^−4^, weight decay 0.01, dropout 0.1, and batch size 32 for 100 epochs.

#### siRNA Efficacy Prediction

This task evaluates siRNA potency across two benchmark datasets. It is a regression task evaluated with AUROC and AUPRC. We adopt the Oligoformer framework and replace the RNA-FM encoder with RNAGenesis. The model is trained using AdamW with a learning rate of 1 × 10^−4^, dropout 0.1, batch size of 64, and trained for 200 epochs.

#### Human UTR Variant Prediction (UTR-P, UTR-Reg)

We consider two subtasks: (1) classifying UTR variants as pathogenic or benign (UTR-P) and (2) predicting up-regulating, down-regulating, or neutral effects (UTR-Reg) based on MPRA measurements. Both are sequence-level classification tasks. Token embeddings from the last RNAGenesis layer are mean-pooled and fed into a linear classifier. UTR-P is treated as binary classification; UTR-Reg as three-class classification. We use cross-entropy loss and evaluate using AUROC and AUPRC. Fine-tuning uses AdamW (learning rate 1 × 10^−4^, weight decay 0.01), batch size 32, and 100 epochs.

#### Aptamer Generation and Evaluation

This task evaluates the generative design of RNA aptamers that mimic SELEX-enriched candidates. We freeze the pretrained RNAGenesis encoder and fine-tune a Q-former, adapted from the ESM-2 architecture^75^, along with a decoder based on ProGen2-small^76^. The model is trained with teacher forcing using AdamW (learning rate 2 × 10^−4^), batch size of 64, and trained for 10 epochs. Evaluation includes sequence edit distance, GC content, minimum free energy (MFE), and secondary structure edit distance.

### More Details of sgRNA Scaffold Design Data

For sgRNA scaffold design, we leverage the SELEX data from Bush, K. et al.^63^ to fine-tune RNAGenesis. The study by Bush, K. et al.^63^ employed a BLADE (Binding and Ligand Activated Directed Evolution) approach, a functional SELEX method, to evolve diverse sgRNA scaffolds from a partially randomized library based on the Streptococcus pyogenes sgRNA sequence. Their selection process involved five rounds with distinct partitioning strategies: rounds 1 and 2 focused on sgRNA binding to Cas9 via nitrocellulose filter binding, round 3 selected for RNP complexes binding to target DNA using biotinylated DNA and streptavidin beads, and rounds 4 and 5 isolated cleavage-capable sgRNAs by employing a TdT-based approach to tag and purify cleaved DNA products. While the BLADE approach successfully identified diverse sgRNA variants, it suffers from inefficiencies inherent to SELEX-based methods, such as the labor-intensive and time-consuming nature of iterative in vitro selection and partitioning, which limits scalability and rapid optimization for specific genomic targets^63^. In RNAGenesis, we combine strong generation capabilities and bio insights to design sgRNAs with enhanced properties.

### More Analysis of Secondary Structural Prediction

Besides benchmark results, we aim to provide further insights into secondary structure predictions. When evaluating prediction results, it is crucial to account for the inherent structural dynamics of RNA. Consequently, predictions that are “close enough” to the true pairings are treated as correct. Following the approach proposed by Mathews et al.^103^, we therefore consider a pairing (*i*, *j*) to be correct if the predicted pair differs by at most one nucleotide in either index. Specifically, if (*i*, *j*) is the true pairing, then (*i* ± 1, *j*) and (*i*, *j* ± 1) are also counted as correct predictions (**soft**). To maintain alignment with Mathews et al.^103^, we also determine the optimal threshold for F1 score calculation based on the results from the validation set. To convert base-pairing probabilities into a valid secondary structure, we implement a simple greedy approach (**greedy**). We iteratively fix the nucleotide pair with the highest pairing probability and then exclude all clashing pairs from further iterations. Throughout this process, we ignore non-canonical base pairs (**NC filtering**) and disallow any pairing that would produce a “sharp” hairpin loop (i.e., the *i*th nucleotide cannot be paired with the *j*th nucleotide if |*i* – *j*| < 4) (**sharp**).

In Figure. S5, we show the results of ablation studies. We observe that sharp+soft+greedy leads to the best performance while ignoring non-canonical base pairs brings a drop in performance. This may be explained by the fact that non-canonical base pairs frequently contribute to RNA stability and folding, especially in complex motifs. By filtering them out, the model loses potentially informative interactions, resulting in predictions that deviate more strongly from the true conformation. Consequently, while the **sharp** and **soft** adjustments effectively capture local structural flexibility, discarding these non-canonical pairings restricts the model’s ability to identify important stabilizing contacts, thus lowering predictive accuracy.

In Figure. S5, we demonstrate the secondary structure prediction examples compared with RNAStructure^104^, which relies on thermodynamic models. Compared to thermodynamic methods, our model can achieve more accurate predictions. At the same time, we have found that our model can predict very well on short sequences, and there is room for improvement on long sequences, which we will further optimize in the future.

## Supplementary Figures

**Figure S1.**
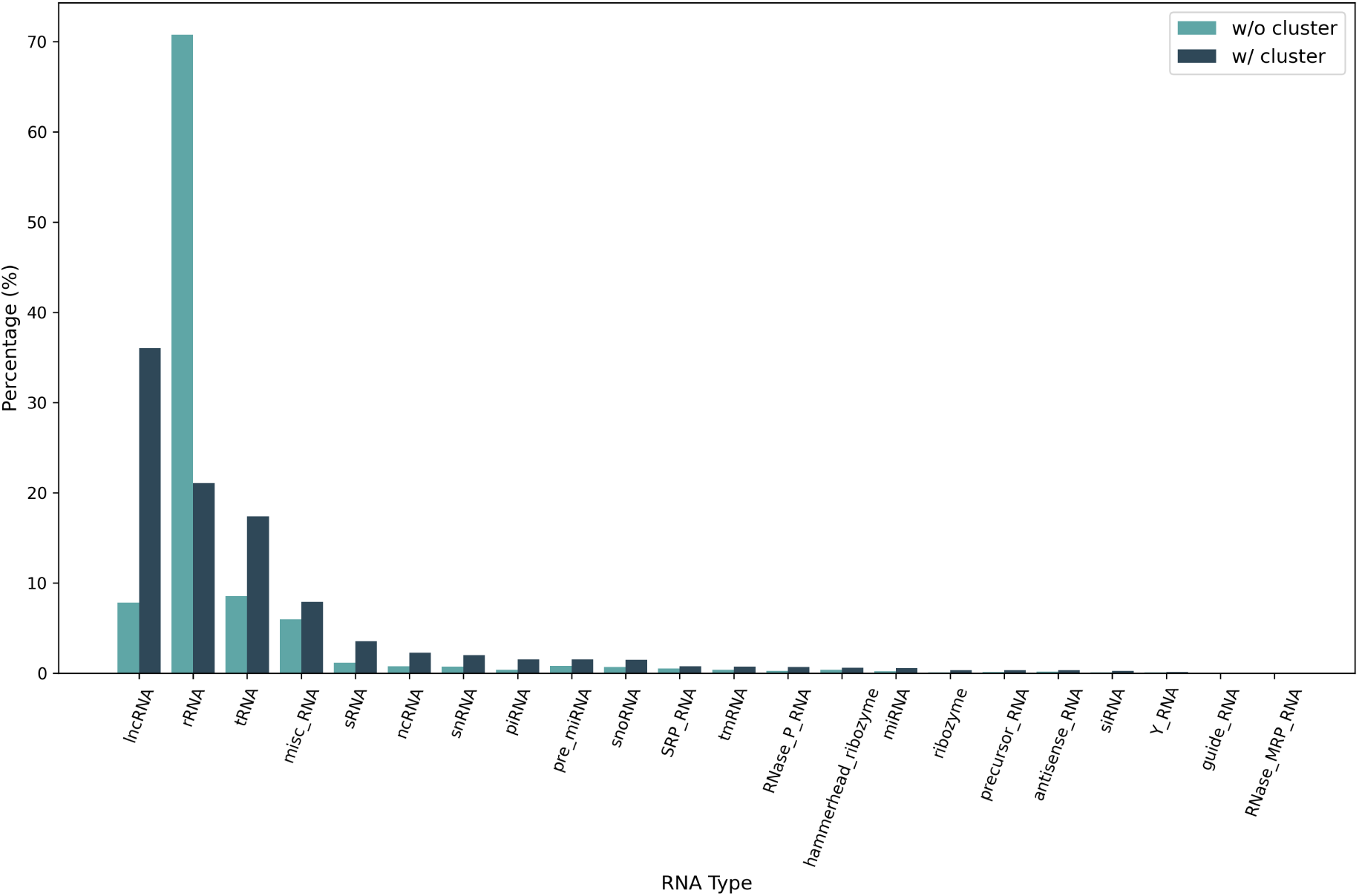
Data distribution comparison between clustered and non-clustered version.

**Figure S2.**
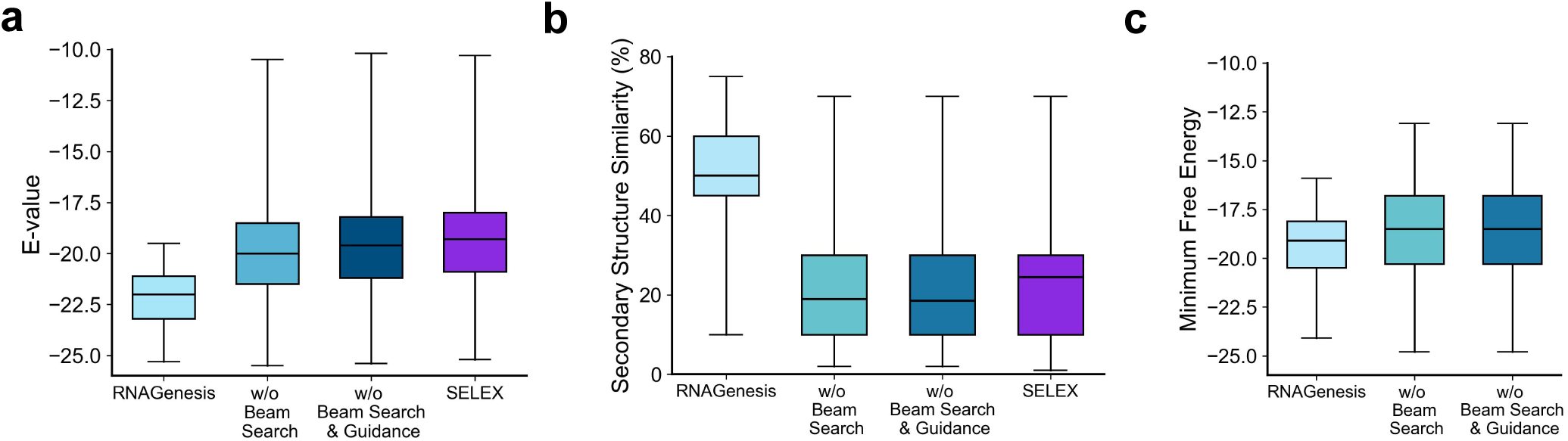
Ablation studies of Beam Search and Guidance for RNAGenesis. The Beam Search and Guidance methods are removed from RNAGenesis generation for ablation to show their contribution. The E-value, secondary structure similarity, and minimum free energy are compared. The corresponding distributions of the SELEX dataset are shown as a reference.

The impact of model scaling on learning efficiency and generalization remains a fundamental question in language model research. In this study, we systematically investigate the scaling laws governing RNAGenesis encoders by training four models with different parameter sizes while maintaining identical training settings. Each model follows a Transformer-based architecture, with parameter growth achieved by scaling hidden dimensions, depth, and attention heads. The smallest model, 40M parameters, has a hidden size of 512, with 8 layers and 8 attention heads. As the model scales up, the 80M parameter variant increases the hidden size to 640, expands to 12 layers, and employs 20 attention heads, allowing for greater representation capacity. The 180M parameter model maintains the same hidden size of 640 but significantly deepens the architecture to 30 layers, retaining 20 attention heads for efficient information integration. Finally, the largest model, 700M parameters, further scales the hidden size to 1280, with 32 layers and 20 attention heads, maximizing both depth and expressiveness. Despite these architectural differences, all models share the same optimization strategy, learning rate schedule, and tokenization process to ensure comparability in the experimental setup. As demonstrated in the Figure. S3, model performance improves consistently with increased parameter count. Larger models exhibit both faster initial convergence and lower cross-entropy loss. These results align with prior findings in protein language models^75, 105^.

**Figure S3.**
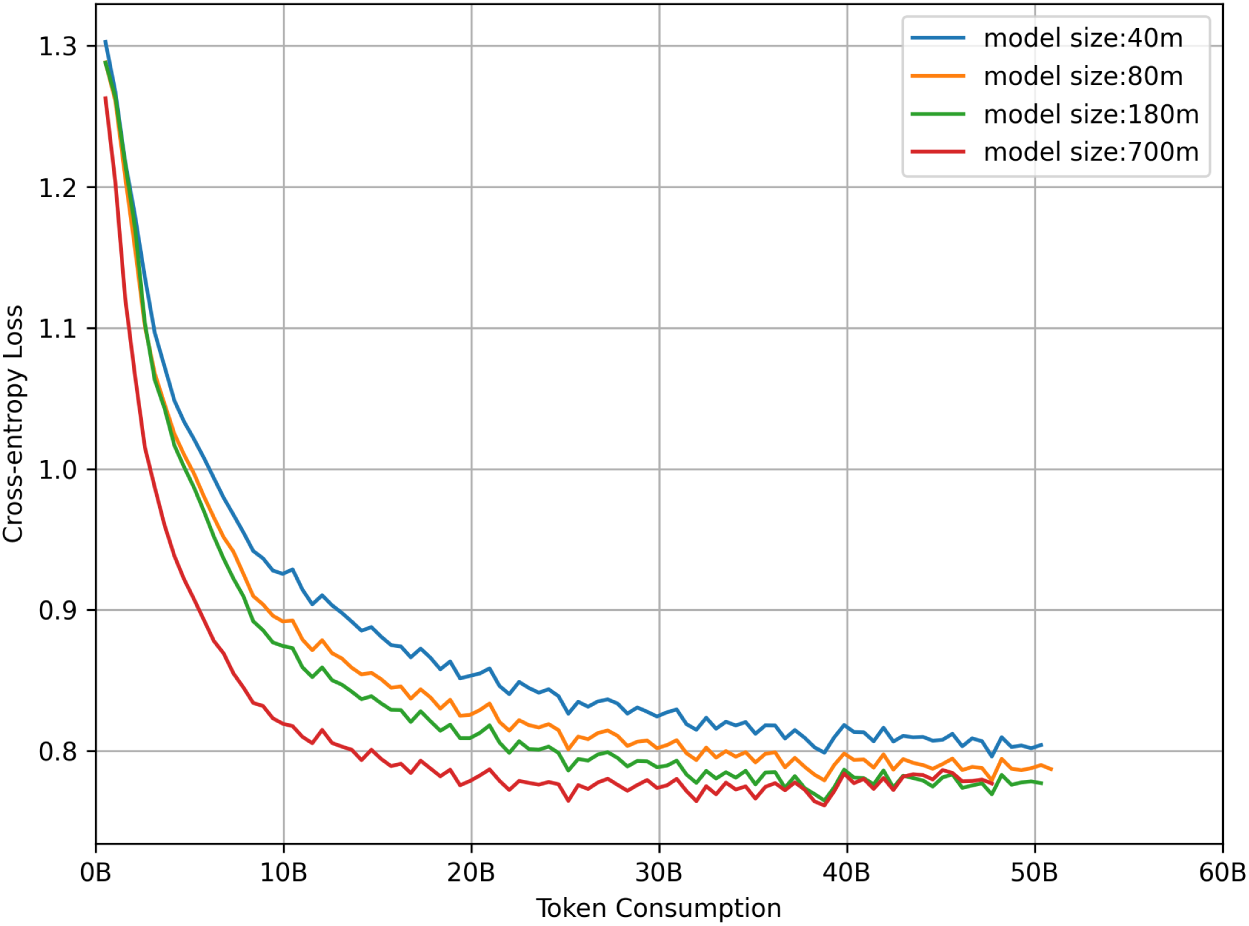
Empirical Study on the Scaling Law of RNAGenesis Encoder.

**Figure S4.**
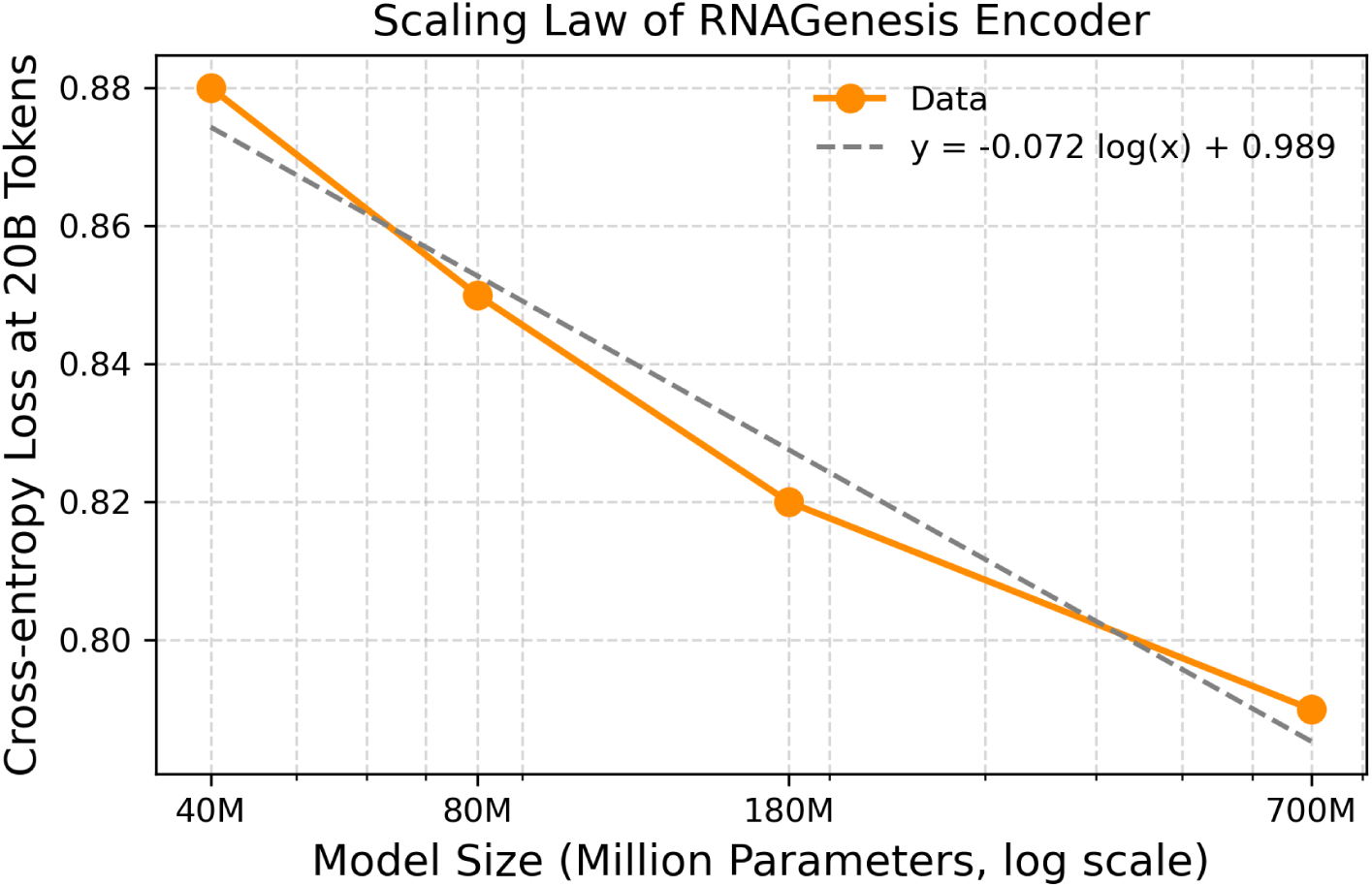
The scaling trend of RNAGenesis encoder between the model size and the model performance (quantified by training cross-entropy loss).

**Figure S5.**
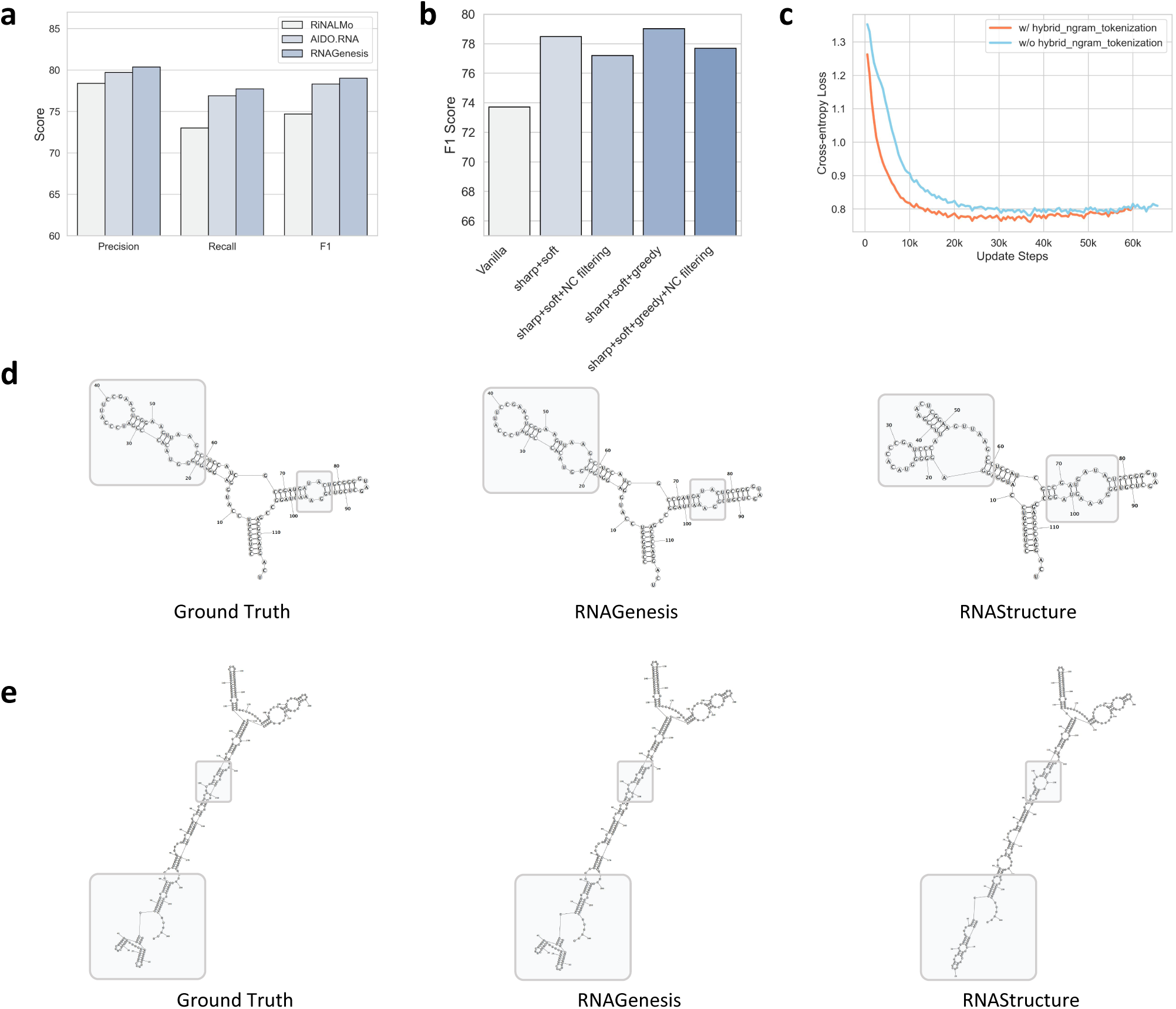
Futher Analysis of RNAGenesis. (**a**) Benchmarking RNAGenesis with two state-of-the-art baselines RiNALMo and AIDO.RNA on Secondary Structure Prediction. Precision, Recall, and F1 scores are reported. The Recall and F1 values of RiNALMo and AIDO.RAN are borrowed from their original papers. The Precision values are then calculated based on Recall and F1. (**b**) Ablation studies of RNAGenesis on Secondary Structure Prediction. The post-processing strategies include disallowing sharp hairpin loop (**sharp**), ignoring non-canonical base pairs (**NC filtering**), greedy approach for pairing (**greedy**), and considering a pairing to be correct if the predicted one differs by at most one nucleotide (**soft**). Besides, the classification threshold was tuned on the validation set. (**c**) Ablation study on the Hybrid N-Gram Tokenization module. The inclusion of the Hybrid N-Gram Tokenization module leads to faster convergence of the pretraining loss and a lower final loss value. (**d** & **e**) Example of secondary structure prediction. The examples show RNAGenesis can achieve nearly perfect prediction results on short sequences, and it also performs well on longer sequences, although the results are slightly inferior. In the future, we will further enhance the prediction capability for long sequences. The gray boxes highlight the RNA regions where RNAGenesis achieves better prediction than RNAStructure.

**Figure S6.**
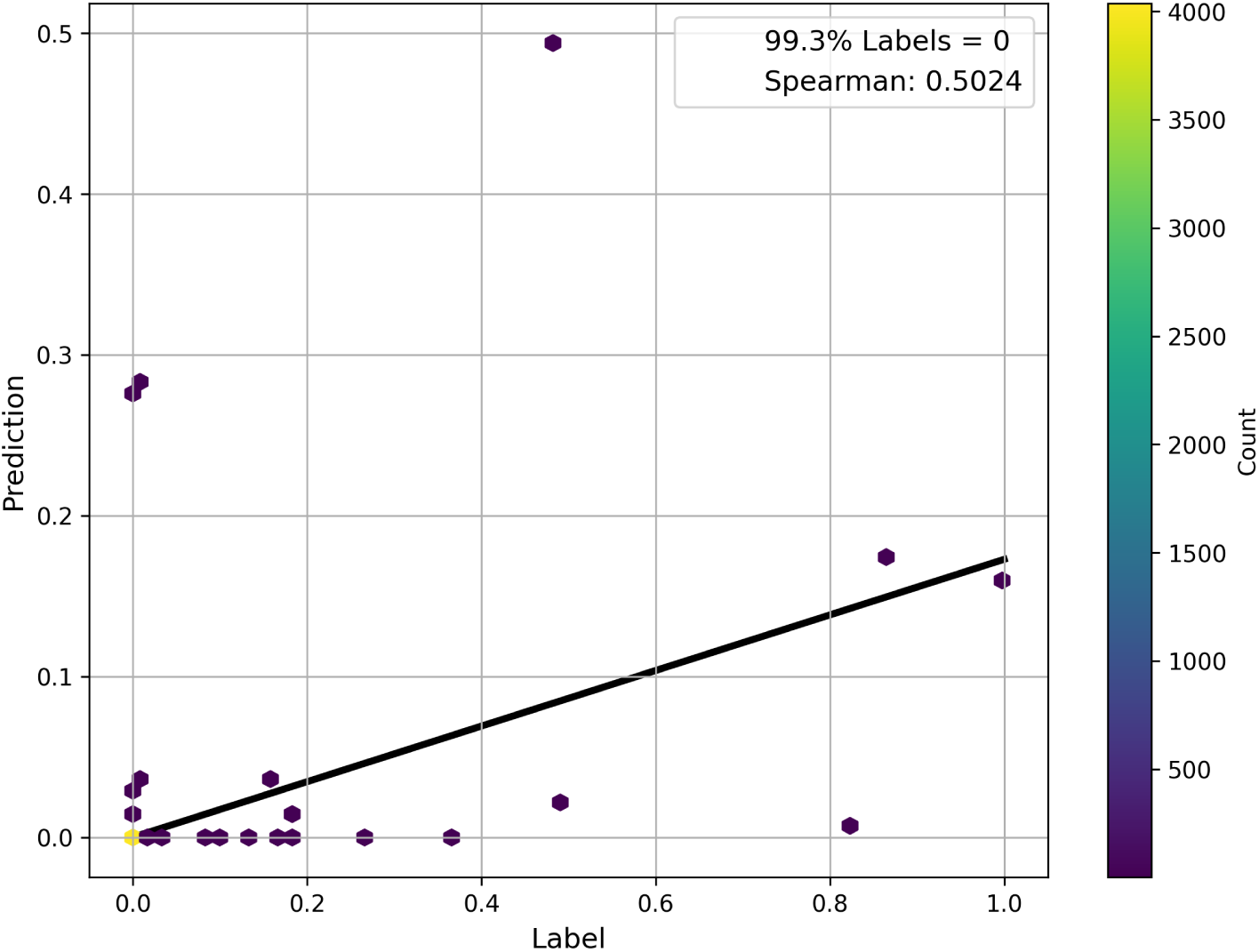
Scatter plots depicting the results of RNAGenesis on CRISPR Off-Target Prediction task.

We observe a severe label imbalance in CRISPR Off-Target benchmark, where over 99% of data points have zero-value labels (Figure. S6). This distribution skew makes the model prone to zero-value predictions, while its predictive performance on non-zero labels critically determines the Spearman correlation coefficient - our primary evaluation metric. As demonstrated in Figure. S6, our model achieves both accurate predictions for non-zero labels and superior Spearman scores, indicating its robustness against label imbalance.

**Figure S7.**
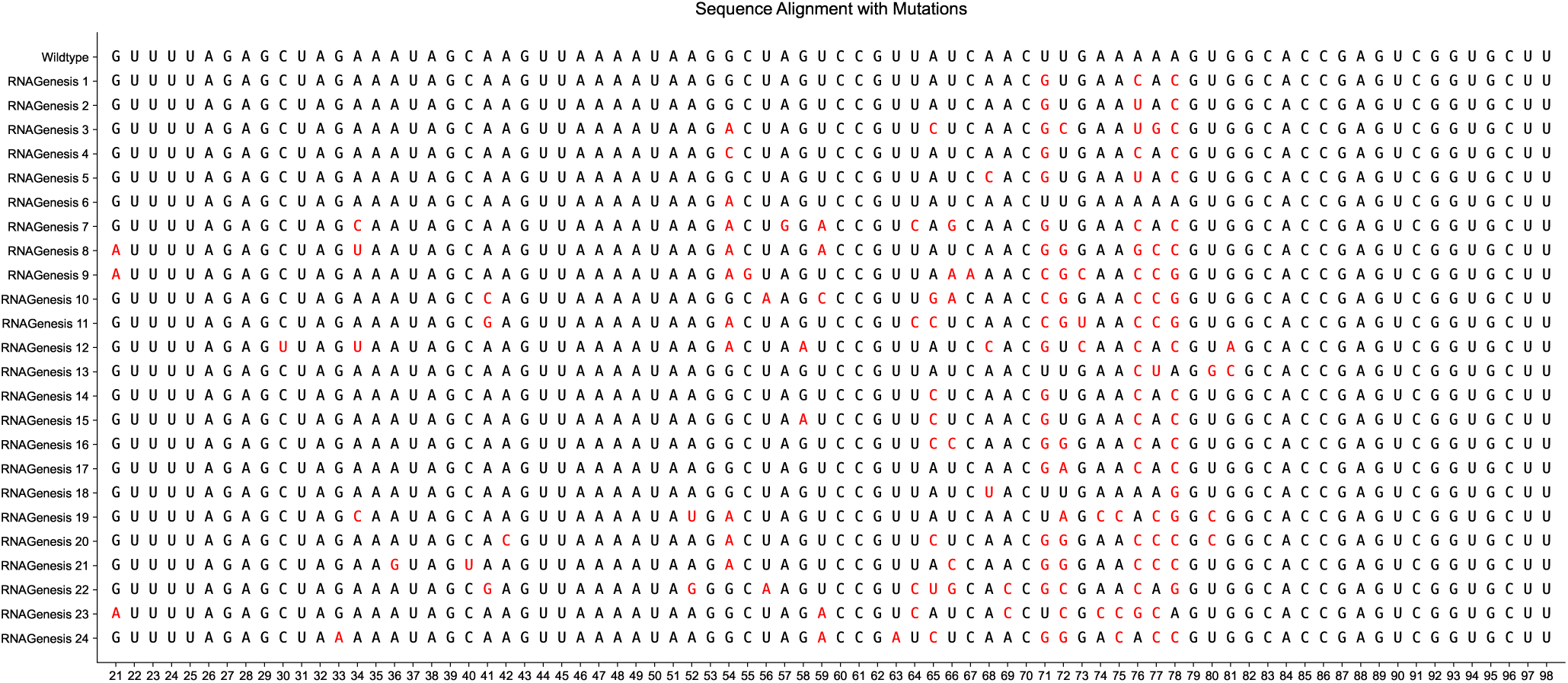
Aligning RNAGenesis-designed sequences with wild-type sgRNA. The mutated nucleic acids are colored red. In Figure. 4, RGen 1-6 correspond to sequences 1,3,14,15,16,24.

**Figure S8.**
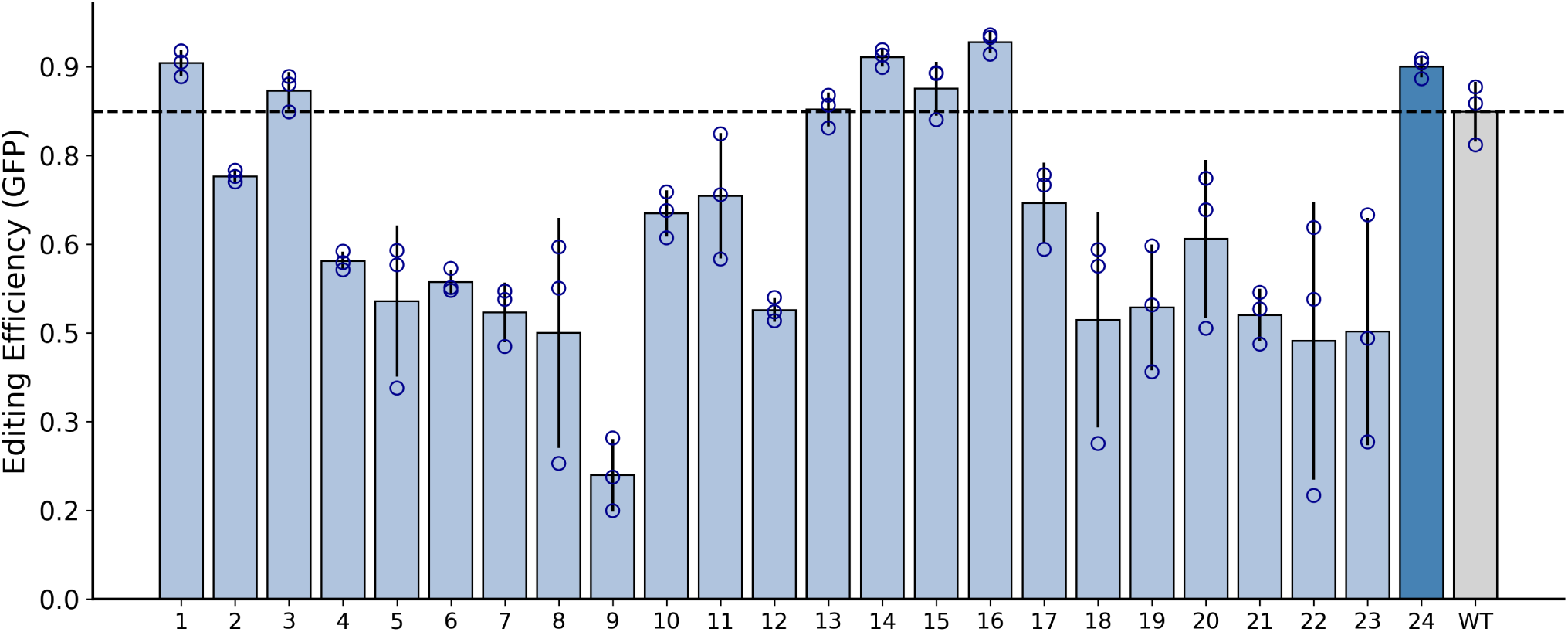
Editing Efficiency for GFP. In Figure. 4, RGen 1-6 correspond to sequences 1,3,14,15,16,24.

**Figure S9.**
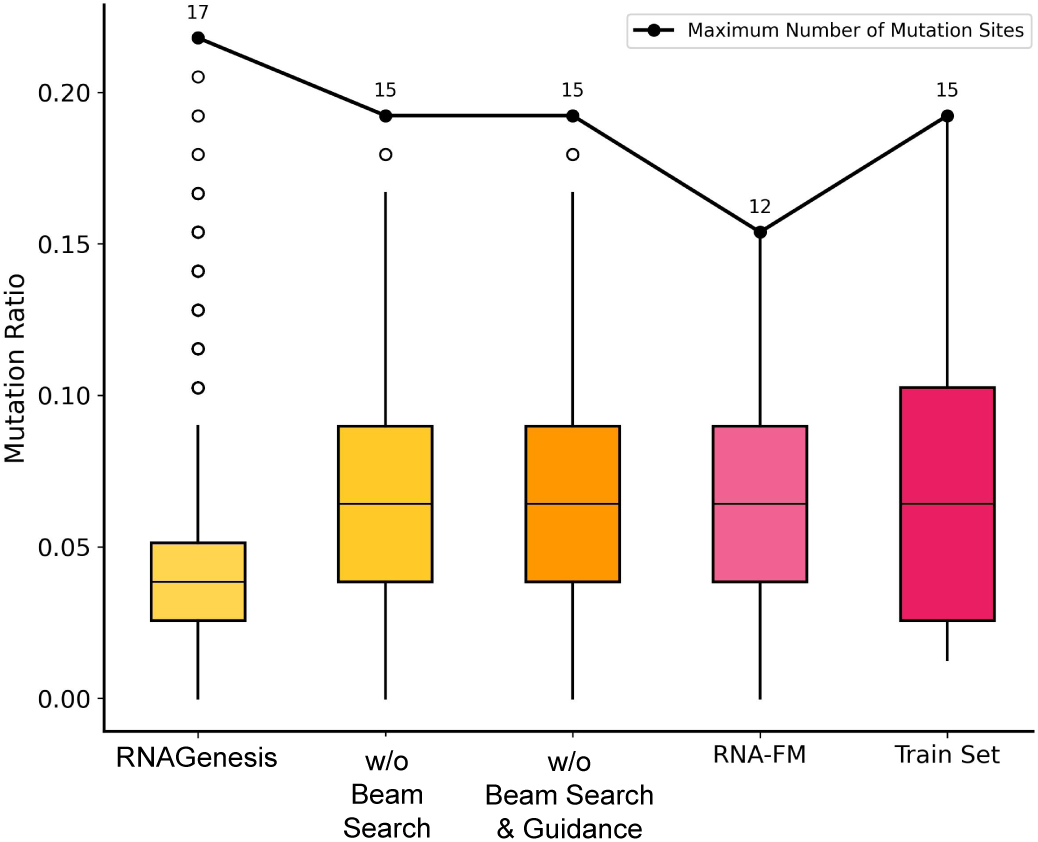
Comparison of mutation frequencies in highly homologous targets using sgRNAs designed by RNAGenesis and other methods. Line chart shows the comparison of maximum mutation sites in homologous targets using different sgRNA design methods.

**Figure S10.**
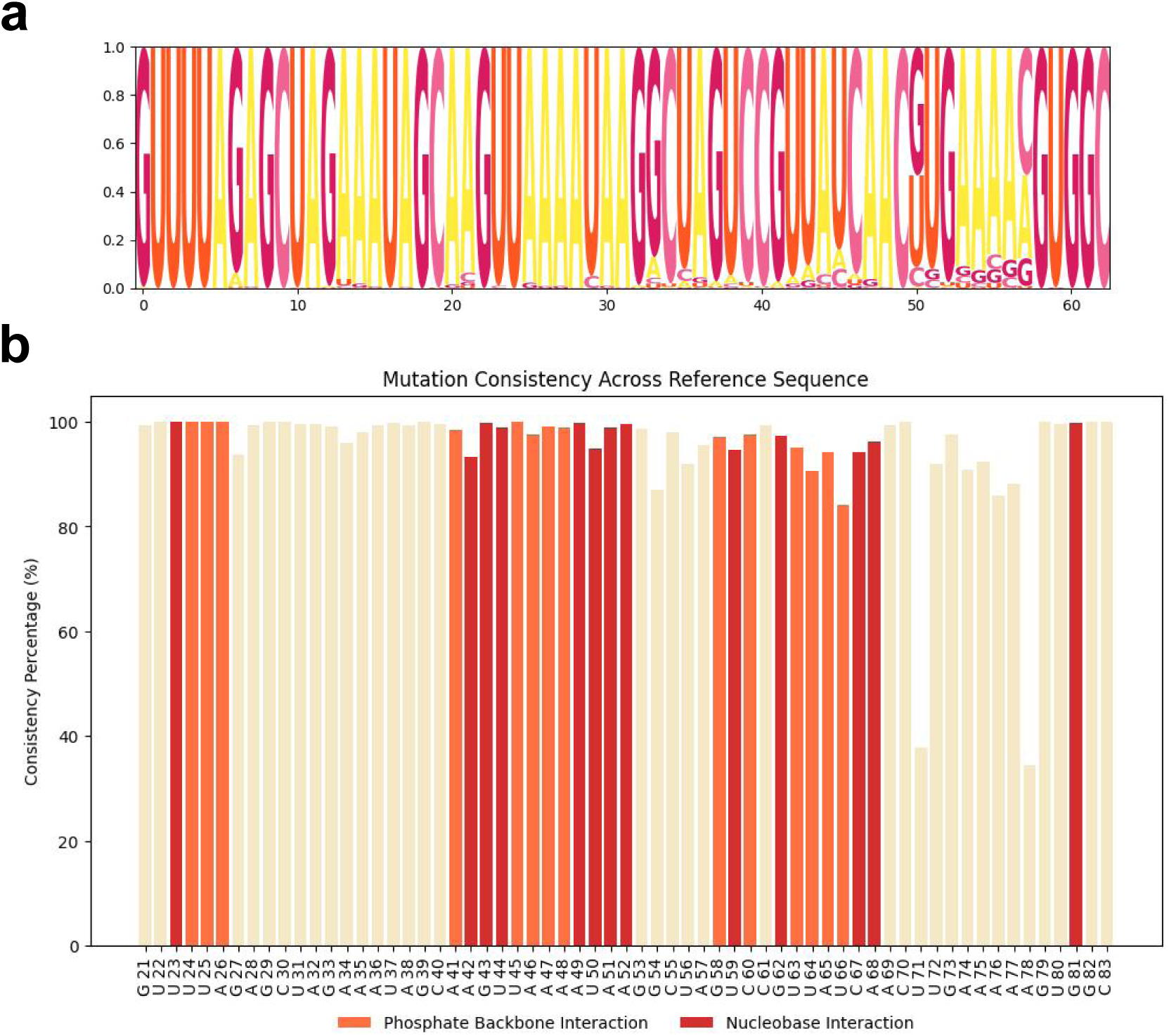
Diversity of generated sequences by RNAGenesis. (**a**) Logo plot of sgRNA sequences generated by RNAGenesis, showing nucleotide diversity and conservation at each position. (**b**) Conservation of each nucleotide position in RNAGenesis-generated sgRNA sequences relative to the wild-type sequence, highlighting differences from the original nucleotide types.

**Figure S11.**
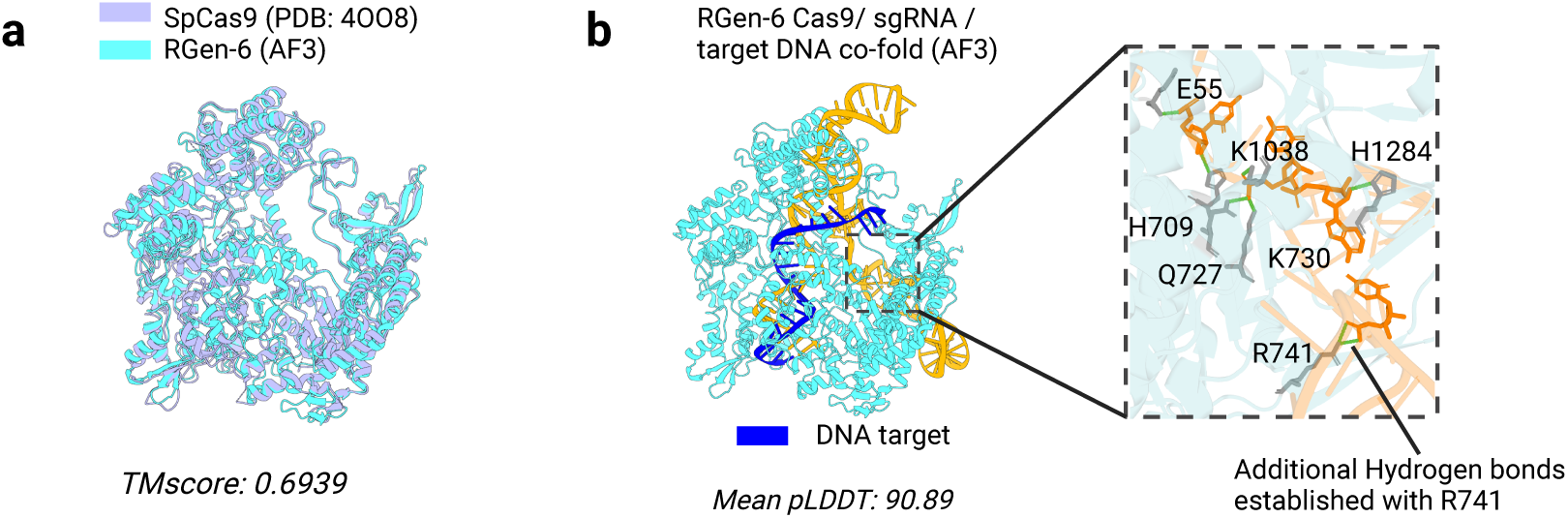
(**a**) AlphaFold3 predicted RGen-6 Cas9 aligned with SpCas9 from PDB 4OO8. (**b**) RNAGenesis Designed sgRNA scaffold establishes more hydrogen bonds with Cas9. The Structure is predicted with AlphaFold3. Hydrogen bonds are detected with MDAnalysis.

**Figure S12.**
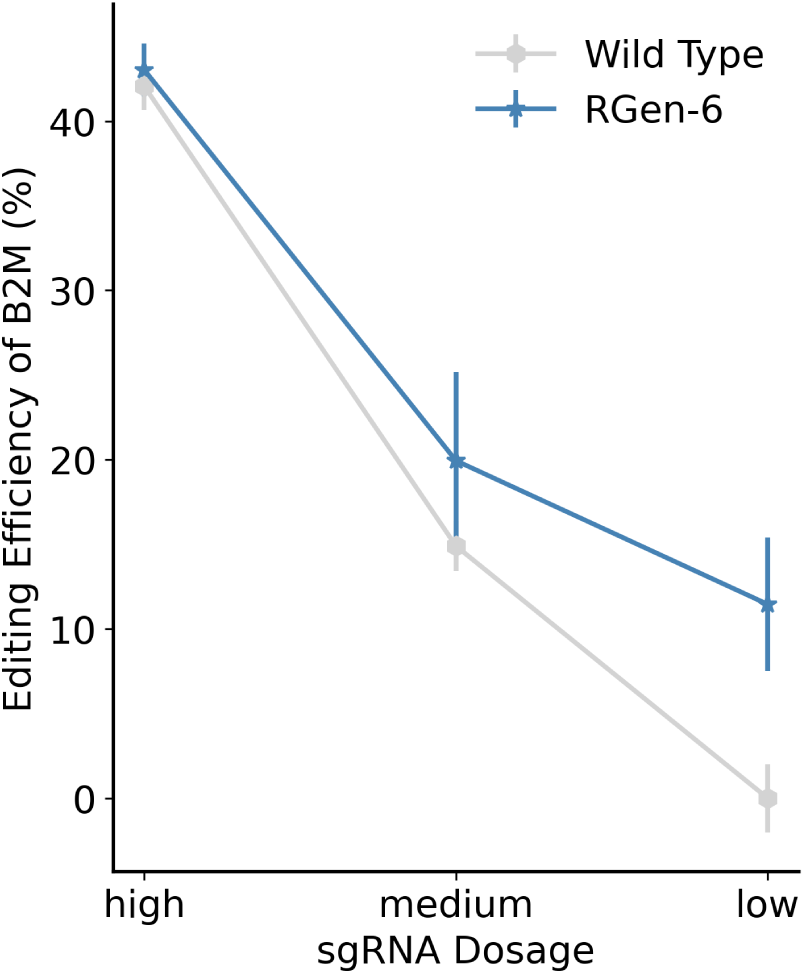
Editing efficiency of RGen-6 targeting B2M compared with wild type across a range of sgRNA dosages (high: 80ng, medium: 40ng, low: 20ng).

**Figure S13.**
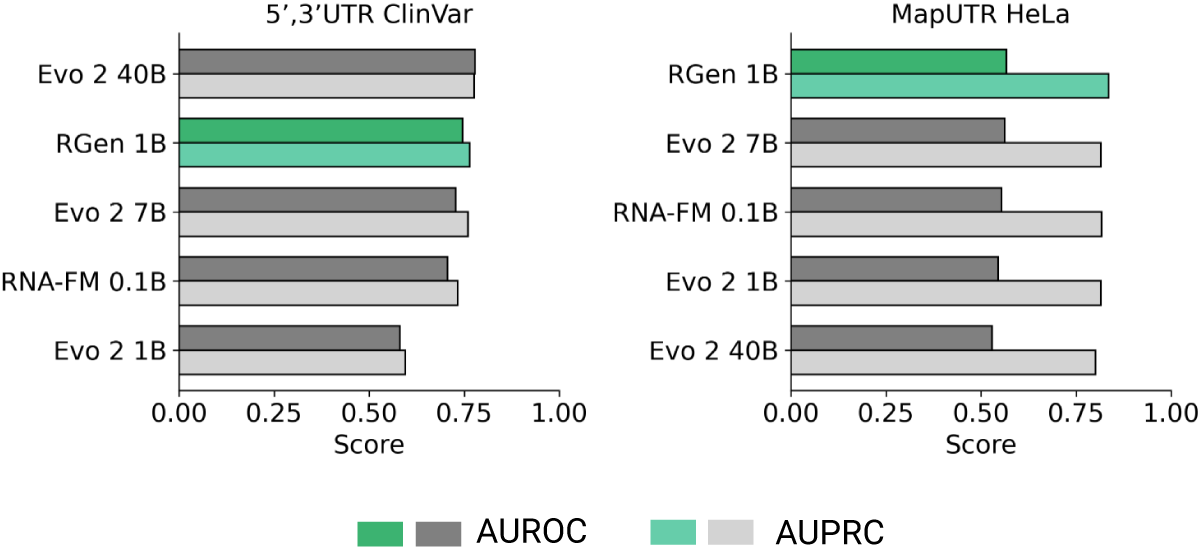
Performance on predicting the effects of human UTR variants. Tasks include classifying variants as pathogenic or benign (*N* = 6,265 for ClinVar datasets) and as up-regulating or down-regulating (*N* = 17,210 for MapUTR HeLa datasets).

**Figure S14.**
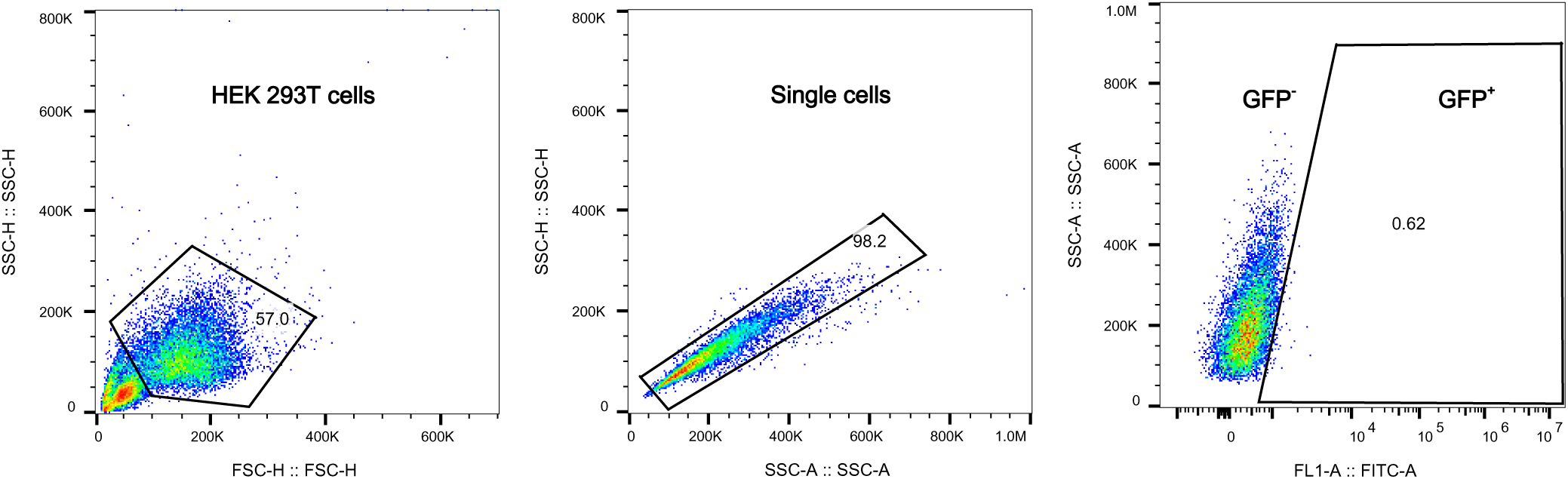
Gating strategies for Base Editing reporter assay.

**Figure S15.**
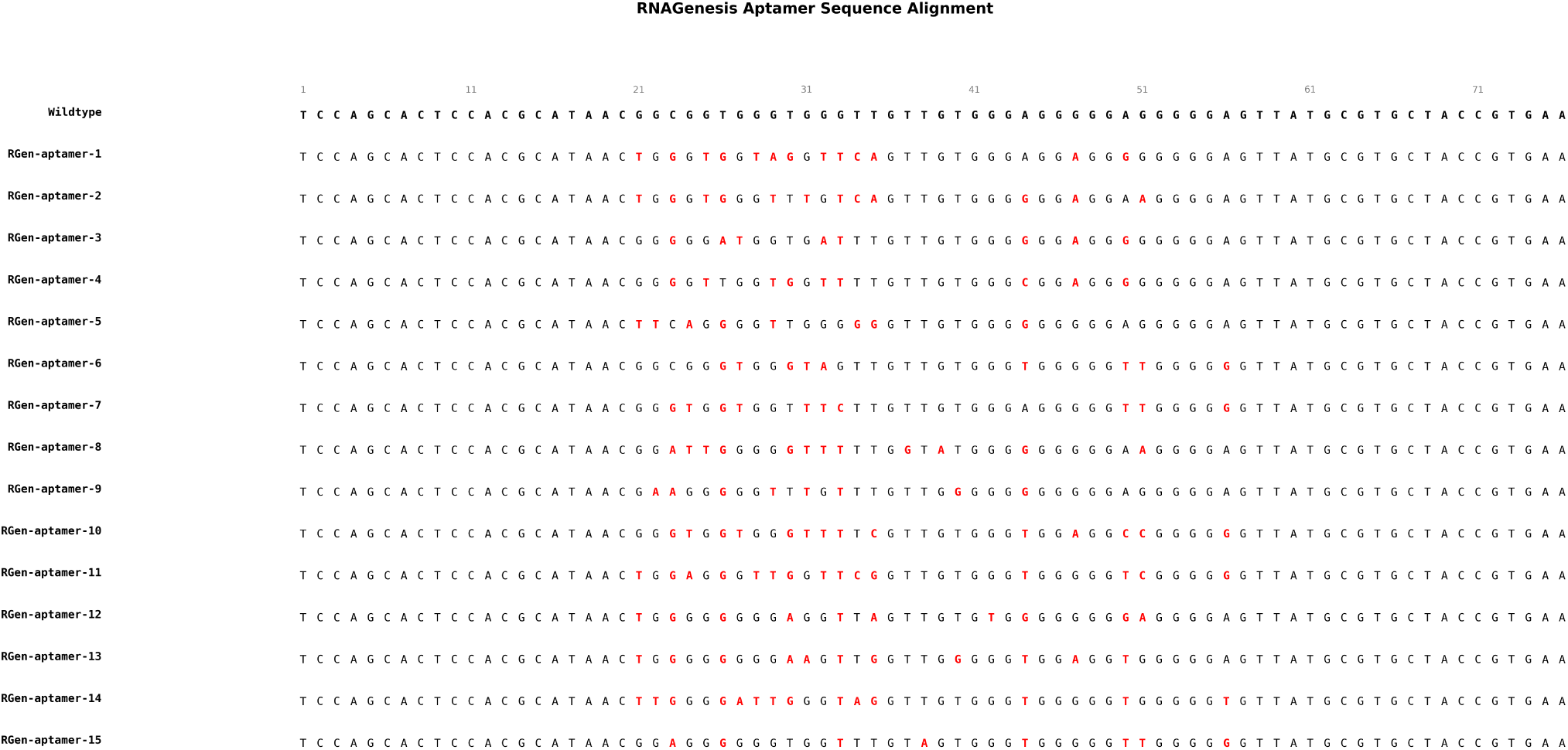
Sequence alignment of RNAGenesis-designed aptamers against the SELEX wildtype. Nucleotide sequences of 15 aptamers generated by RNAGenesis are aligned to a SELEX-derived wildtype reference. Positions with mutations relative to the wildtype are highlighted in red.

**Figure S16.**
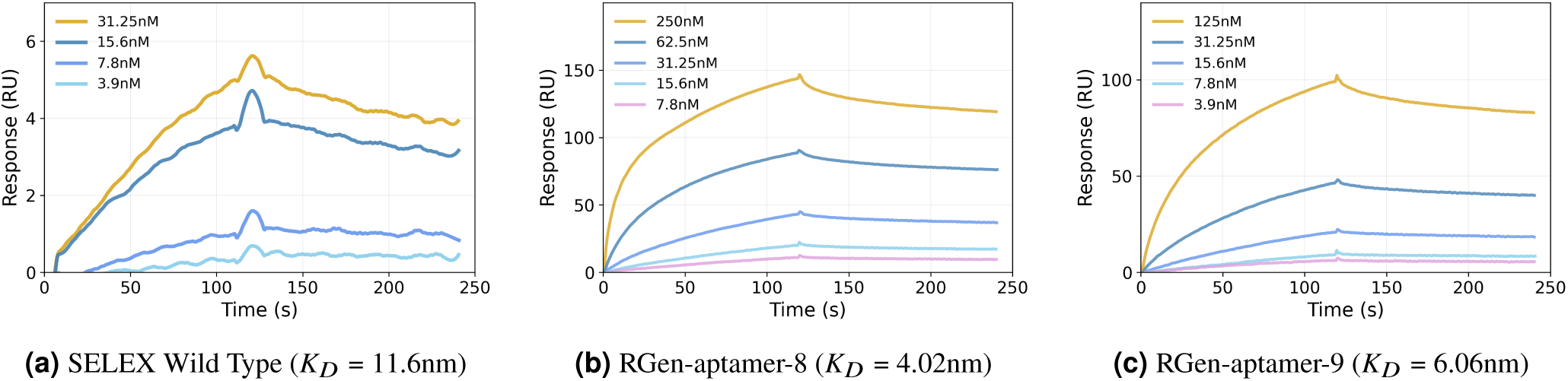
Surface plasmon resonance (SPR) sensorgrams comparing the binding affinities of SELEX and RNAGenesis-designed aptamers. (a) SELEX-derived wildtype aptamer, (b) RGen-aptamer-8, and (c) RGen-aptamer-9 were evaluated for binding to IGFBP3 using SPR. Each curve represents a different aptamer concentration, with response units (RU) plotted over time. Both RGen-aptamer-8 and RGenaptamer-9 demonstrate stronger binding responses than the SELEX wildtype, indicating improved affinity.

**Figure S17.**
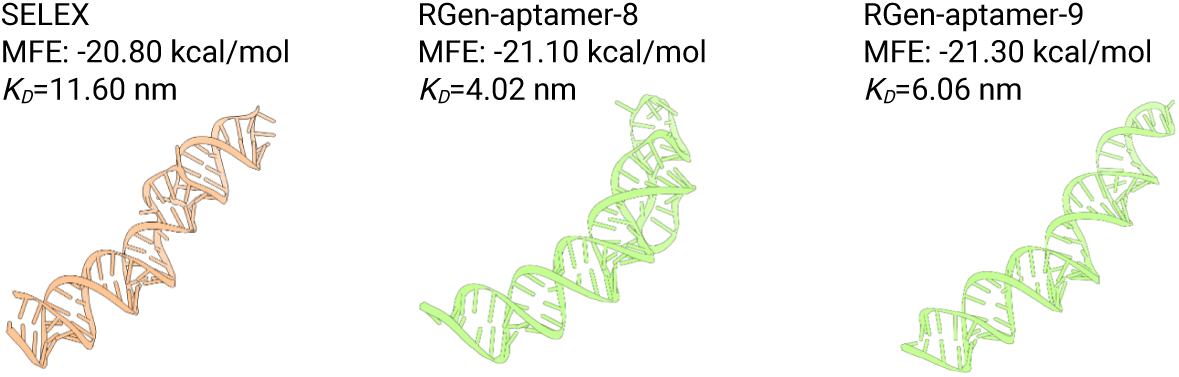
AlphaFold3 predicted structures of wildtype aptamer and RNAGenesis designed ones. MFE and *K*_*D*_ are annotated.

## Supplementary Tabls

**Table S1.**
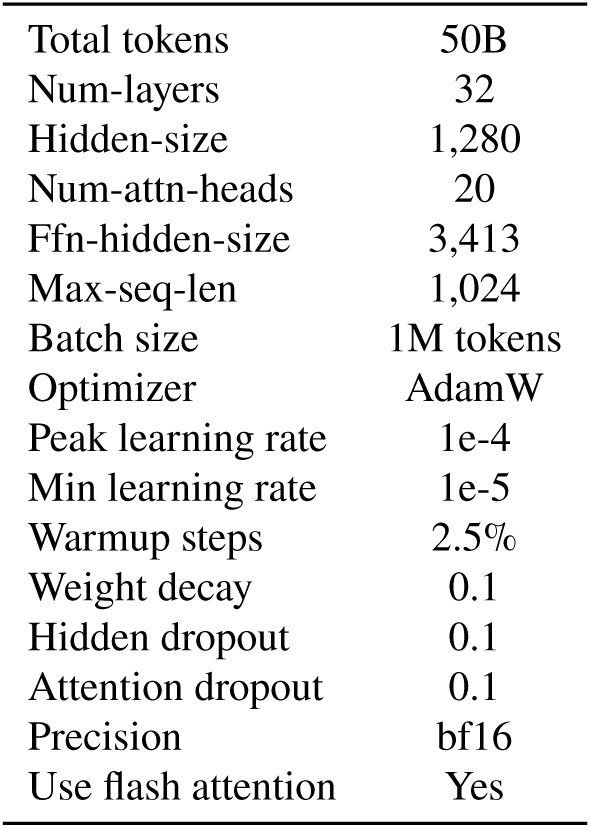
Hyperparameters for pre-training RNAGenesis encoder module.

|  |  |
| --- | --- |
| Total tokens | 50B |
| Num-layers | 32 |
| Hidden-size | 1,280 |
| Num-attn-heads | 20 |
| Ffn-hidden-size | 3,413 |
| Max-seq-len | 1,024 |
| Batch size | 1M tokens |
| Optimizer | AdamW |
| Peak learning rate | 1e-4 |
| Min learning rate | 1e-5 |
| Warmup steps | 2.5% |
| Weight decay | 0.1 |
| Hidden dropout | 0.1 |
| Attention dropout | 0.1 |
| Precision | bf16 |
| Use flash attention | Yes |

**Table S2.** Related work of RNA foundation models.

| FM | Architecture | Model size | Pre-training dataset | Tokenization | Pre-training objective |
| --- | --- | --- | --- | --- | --- |
| RNABERT<sup>5</sup> | encoder-only transformer | <10M | RNAcentral (76,237 human-derived small ncRNA sequences) & Rfam alignment | base | MLM(15%) + others |
| RNA-FM<sup>6</sup> | encoder-only transformer | 100M | RNAcentral v19 (23M ncRNA sequences) | base | MLM(15%) |
| RNA-MSM<sup>106</sup> | MSA transformer | ~200M | Rfam v14.7 | base | MLM(20%) |
| Uni-RNA<sup>11</sup> | encoder-only transformer | <400M | RNAcentral & nt & Genome Warehouse (1B potential RNA sequences) | base | MLM |
| SpliceBERT<sup>8</sup> | encoder-only transformer | 20M | 2M pre-mRNA sequences from UCSC genome browser | base | MLM(15%) |
| CodonBERT<sup>14</sup> | encoder-only transformer | ~100M | NCBI (10M mRNA coding sequences) | codon | MLM(15%) + others |
| 3UTRBERT<sup>9</sup> | encoder-only transformer | ~100M | 86k 3'UTRs curated from GRCh38.p13, Release 40 | k-mer | MLM(15%) |
| UTR-LM<sup>7</sup> | encoder-only transformer | <10M | Ensembl & eGFP & mCherry & Cao (700K 5' UTRs) | base | MLM + others |
| ATOM-1<sup>107</sup> | encoder-decoder transformer | unknown | chemical mapping data | base | unknown |
| RiNALMo<sup>10</sup> | encoder-only transformer | 650M | RNAcentral & Rfam & nt & Ensembl (36M unique ncRNA sequences) | base | MLM |
| CaLM<sup>83</sup> | encoder-only transformer | 86M | European Nucleotide Archive (9M cDNA sequences) | codon | MLM(25%) |
| RNAErnie<sup>13</sup> | encoder-only transformer | ~100M | RNAcentral (23M ncRNA sequences) | base | MLM(15%) + others |
| AIDO.RNA<sup>11</sup> | encoder-only transformer | 1.6B | RNAcentral v24.0 (42M ncRNA sequences) | base | MLM(15%) |
| RNAGenesis (ours) | encoder-decoder transformer | ~1B | RNAcentral v24.0 (42M ncRNA sequences) | Hybrid N-Gram | MLM(15%) |

**Table S3.** Ablation study on Hybrid N-Gram Tokenization for key sequence understanding tasks.

| Model | CMP | MRL | Modif | CRI-Off |
| --- | --- | --- | --- | --- |
| RNAGenesis | $72.10 \pm 0.23$ | $85.83 \pm 0.05$ | $94.98 \pm 0.10$ | $44.93 \pm 3.77$ |
| RNAGenesis w/o Hybrid N-Gram Tokenization | $70.48 \pm 0.69$ | $85.38 \pm 0.27$ | $94.9 \pm 0.18$ | $34.02 \pm 9.86$ |

## Notes

### Competing Interest Statement

The authors have declared no competing interest.

### Summary of Updates

correct the typo in figure 3, rearrange the supplementary

## References

1 Lu, T. X. & Rothenberg, M. E. Microrna. Journal of allergy and clinical immunology 141, 1202–1207 (2018).

2 Mercer, T. R., Dinger, M. E. & Mattick, J. S. Long non-coding rnas: insights into functions. Nature reviews genetics 10, 155–159 (2009).

3 Hu, B. et al. Therapeutic sirna: state of the art. Signal transduction and targeted therapy 5, 101 (2020).

4 Cong, L. et al. Multiplex genome engineering using crispr/cas systems. Science 339, 819–823 (2013).

5 Akiyama, M. & Sakakibara, Y. Informative rna base embedding for rna structural alignment and clustering by deep representation learning. NAR genomics and bioinformatics 4, lqac012 (2022).

6 Chen, J. et al. Interpretable rna foundation model from unannotated data for highly accurate rna structure and function predictions. bioRxiv 2022–08 (2022).

7 Chu, Y. et al. A 5’ utr language model for decoding untranslated regions of mrna and function predictions. Nature Machine Intelligence 1–12 (2024).

8. Chen, K., et al. Self-supervised learning on millions of pre-mrna sequences improves sequence-based rna splicing prediction. bioRxiv (2023). URL https://www.biorxiv.org/content/early/2023/05/09/2023.01.31.526427. https://www.biorxiv.org/content/early/2023/05/09/2023.01.31.526427.full.pdf.

9 Yang, Y. et al. Deciphering 3’ utr mediated gene regulation using interpretable deep representation learning. bioRxiv (2023). URL https://www.biorxiv.org/content/early/2023/09/12/2023.09.08.556883. https://www.biorxiv.org/content/early/2023/09/12/2023.09.08.556883.full.pdf.

10 Penić, R. J., Vlašić, T., Huber, R. G., Wan, Y. & Šikić, M. Rinalmo: General-purpose rna language models can generalize well on structure prediction tasks. arXiv preprint arXiv:2403.00043 (2024).

11 Zou, S. et al. A large-scale foundation model for rna function and structure prediction. bioRxiv 2024–11 (2024).

12 Rnacentral 2021: secondary structure integration, improved sequence search and new member databases. Nucleic acids research 49, D212–D220 (2021).

13 Wang, N. et al. Multi-purpose rna language modelling with motif-aware pretraining and type-guided fine-tuning. Nature Machine Intelligence 1–10 (2024).

14 Li, S. et al. Codonbert: Large language models for mrna design and optimization. bioRxiv 2023–09 (2023).

15. Ren, Y., et al. Beacon: Benchmark for comprehensive rna tasks and language models. NeurIPS (2024).

16 Brixi, G. et al. Genome modeling and design across all domains of life with evo 2. BioRxiv 2025–02 (2025).

17 Wong, F. et al. Deep generative design of rna aptamers using structural predictions. Nature Computational Science 1–11 (2024).

18 Porto, E. M., Komor, A. C., Slaymaker, I. M. & Yeo, G. W. Base editing: advances and therapeutic opportunities. Nature Reviews Drug Discovery 19, 839–859 (2020).

19 Scholefield, J. & Harrison, P. T. Prime editing–an update on the field. Gene Therapy 28, 396–401 (2021).

20 Abramson, J. et al. Accurate structure prediction of biomolecular interactions with alphafold 3. Nature 630, 493–500 (2024).

21 Wohlwend, J. et al. Boltz-1: Democratizing biomolecular interaction modeling. bioRxiv 2024–11 (2024).

22. Huang, K., et al. Latent diffusion models for controllable rna sequence generation. arXiv preprint arXiv:2409.09828 (2024).

23. Li, J., Li, D., Savarese, S. & Hoi, S. Blip-2: Bootstrapping language-image pre-training with frozen image encoders and large language models. In International conference on machine learning, 19730–19742 (PMLR, 2023).

24 Anand, R. et al. Rna-frameflow: Flow matching for de novo 3d rna backbone design. arXiv preprint arXiv:2406.13839 (2024).

25 Shi, M. et al. The evolutionary history of vertebrate rna viruses. Nature 556, 197–202 (2018).

26. Huguet, G., et al. Sequence-augmented se (3)-flow matching for conditional protein backbone generation. arXiv preprint arXiv:2405.20313 (2024).

27 Hu, E. J. et al. Lora: Low-rank adaptation of large language models. ICLR 1, 3 (2022).

28. Jing, B., Eismann, S., Suriana, P., Townshend, R. J. & Dror, R. Learning from protein structure with geometric vector perceptrons. arXiv preprint arXiv:2009.01411 (2020).

29 Vaswani, A. Attention is all you need. Advances in Neural Information Processing Systems (2017).

30. Hsu, C. et al. Learning inverse folding from millions of predicted structures. In International conference on machine learning, 8946–8970 (PMLR, 2022).

31. Runge, F., Stoll, D., Falkner, S. & Hutter, F. Learning to design rna. arXiv preprint arXiv:1812.11951 (2018).

32 Das, R. et al. Assessment of three-dimensional rna structure prediction in casp15. Proteins: Structure, Function, and Bioinformatics 91, 1747–1770 (2023).

33 Joshi, C. K. et al. grnade: Geometric deep learning for 3d rna inverse design. bioRxiv 2024–03 (2025).

34. Tan, C., et al. Rdesign: hierarchical data-efficient representation learning for tertiary structure-based rna design. arXiv preprint arXiv:2301.10774 (2023).

35 Rubio-Largo, A., Lozano-Garćıa, N., Granado-Criado, J. M. & Vega-Rodŕıguez, M. A. Solving the rna inverse folding problem through target structure decomposition and multiobjective evolutionary computation. Applied Soft Computing 147, 110779 (2023).

36 Sussman, J. L. et al. Protein data bank (pdb): database of three-dimensional structural information of biological macromolecules. Acta Crystallographica Section D: Biological Crystallography 54, 1078–1084 (1998).

37 Mortimer, S. A., Kidwell, M. A. & Doudna, J. A. Insights into rna structure and function from genome-wide studies. Nature Reviews Genetics 15, 469–479 (2014).

38 Adamczyk, B., Antczak, M. & Szachniuk, M. Rnasolo: a repository of cleaned pdb-derived rna 3d structures. Bioinformatics 38, 3668–3670 (2022).

39 Shen, T. et al. Accurate rna 3d structure prediction using a language model-based deep learning approach. Nature Methods 1–12 (2024).

40 Watkins, A. M., Rangan, R. & Das, R. Farfar2: improved de novo rosetta prediction of complex global rna folds. Structure 28, 963–976 (2020).

41 Li, Y. et al. Integrating end-to-end learning with deep geometrical potentials for ab initio rna structure prediction. Nature Communications 14, 5745 (2023).

42 Magnus, M. et al. Rna-puzzles toolkit: a computational resource of rna 3d structure benchmark datasets, structure manipulation, and evaluation tools. Nucleic acids research 48, 576–588 (2020).

43 Wang, F., Zuroske, T., Watts, J. K. et al. Rna therapeutics on the rise. Nat Rev Drug Discov 19, 441–442 (2020).

44 Rinaldi, C. & Wood, M. J. Antisense oligonucleotides: the next frontier for treatment of neurological disorders. Nature Reviews Neurology 14, 9–21 (2018).

45 Hwang, G. et al. Asoptimizer: Optimizing antisense oligonucleotides through deep learning for ido1 gene regulation. Molecular Therapy-Nucleic Acids 35 (2024).

46 Zhang, G. et al. Identifying circular rna and predicting its regulatory interactions by machine learning. Frontiers in genetics 11, 655 (2020).

47 Bai, Y., Zhong, H., Wang, T. & Lu, Z. J. Oligoformer: an accurate and robust prediction method for sirna design. Bioinformatics 40, btae577 (2024).

48 Zhao, C. et al. Ilgbmsh: an interpretable classification model for the shrna target prediction with ensemble learning algorithm. Briefings in Bioinformatics 23, bbac429 (2022).

49 Ter Brake, O., et al. Lentiviral vector design for multiple shrna expression and durable hiv-1 inhibition. Molecular Therapy 16, 557–564 (2008).

50 Aquino-Jarquin, G. & Toscano-Garibay, J. D. Rna aptamer evolution: two decades of selection. International journal of molecular sciences 12, 9155–9171 (2011).

51 Zhu, Y., Zhu, L., Wang, X. & Jin, H. RNA-based therapeutics: an overview and prospectus. Cell Death Dis. 13, 644 (2022).

52. Zhang, Y., et al. Decoding the rna interactome by ultragen (2024).

53 Zhang, Y. et al. Single-step discovery of high-affinity rna ligands by ultraselex. Nature Chemical Biology 1–9 (2025).

54 Wang, Z. et al. Aptadiff: de novo design and optimization of aptamers based on diffusion models. Briefings in Bioinformatics 25, bbae517 (2024). URL 10.1093/bib/bbae517. https://academic.oup.com/bib/article-pdf/25/6/bbae517/59923079/bbae517.pdf.

55 Lofqvist, C. et al. Igfbp3 suppresses retinopathy through suppression of oxygen-induced vessel loss and promotion of vascular regrowth. Proceedings of the National Academy of Sciences 104, 10589–10594 (2007).

56 Dueñas Rey, A., et al. Combining a prioritization strategy and functional studies nominates 5’utr variants underlying inherited retinal disease. Genome Medicine 16, 7 (2024).

57 Li, G., Wu, J. & Wang, X. Predicting functional utr variants by integrating region-specific features. Briefings in Bioinformatics 25, bbae248 (2024).

58 Fu, T. et al. Massively parallel screen uncovers many rare 3 utr variants regulating mrna abundance of cancer driver genes. Nature Communications 15, 3335 (2024).

59 Doudna, J. A. The promise and challenge of therapeutic genome editing. Nature 578, 229–236 (2020).

60 Jinek, M. et al. A programmable dual-rna–guided dna endonuclease in adaptive bacterial immunity. science 337, 816–821 (2012).

61 Workman, R. E. et al. A natural single-guide rna repurposes cas9 to autoregulate crispr-cas expression. Cell 184, 675–688 (2021).

62 Mali, P. et al. Rna-guided human genome engineering via cas9. Science 339, 823–826 (2013).

63 Bush, K. et al. Utilizing directed evolution to interrogate and optimize crispr/cas guide rna scaffolds. Cell chemical biology 30, 879–892 (2023).

64. Guo, Y., Yuan, H., Yang, Y., Chen, M. & Wang, M. Gradient guidance for diffusion models: An optimization perspective. NeurIPS (2024).

65 Lorenz, R. et al. Viennarna package 2.0. Algorithms for molecular biology 6, 1–14 (2011).

66 Michaud-Agrawal, N., Denning, E. J., Woolf, T. B. & Beckstein, O. Mdanalysis: a toolkit for the analysis of molecular dynamics simulations. Journal of computational chemistry 32, 2319–2327 (2011).

67 Yang, M. et al. Uni-gbsa: an open-source and web-based automatic workflow to perform mm/gb (pb) sa calculations for virtual screening. Briefings in Bioinformatics 24, bbad218 (2023).

68 Crick, F. Central dogma of molecular biology. Nature 227, 561–563 (1970).

69 Su, J. et al. Roformer: Enhanced transformer with rotary position embedding. Neurocomputing 568, 127063 (2024).

70 Dao, T., Fu, D., Ermon, S., Rudra, A. & Ré, C. Flashattention: Fast and memory-efficient exact attention with io-awareness. Advances in Neural Information Processing Systems 35, 16344–16359 (2022).

71. Shazeer, N. Glu variants improve transformer. arXiv preprint arXiv:2002.05202 (2020).

72. Ba, J. L. Layer normalization. arXiv preprint arXiv:1607.06450 (2016).

73 Ho, J., Jain, A. & Abbeel, P. Denoising diffusion probabilistic models. Advances in neural information processing systems 33, 6840–6851 (2020).

74 Li, X., Thickstun, J., Gulrajani, I., Liang, P. S. & Hashimoto, T. B. Diffusion-lm improves controllable text generation. Advances in Neural Information Processing Systems 35, 4328–4343 (2022).

75 Lin, Z. et al. Evolutionary-scale prediction of atomic-level protein structure with a language model. Science 379, 1123–1130 (2023).

76 Nijkamp, E., Ruffolo, J. A., Weinstein, E. N., Naik, N. & Madani, A. Progen2: exploring the boundaries of protein language models. Cell systems 14, 968–978 (2023).

77 Dhariwal, P. & Nichol, A. Diffusion models beat gans on image synthesis. Advances in neural information processing systems 34, 8780–8794 (2021).

78 Guo, Y., Yang, Y., Yuan, H. & Wang, M. Training-free guidance beyond differentiability: Scalable path steering with tree search in diffusion and flow models. arXiv preprint arXiv:2502.11420 (2025).

79. Oshima, Y., Suzuki, M., Matsuo, Y. & Furuta, H. Inference-time text-to-video alignment with diffusion latent beam search. arXiv preprint arXiv:2501.19252 (2025).

80. Houlsby, N. et al. Parameter-efficient transfer learning for nlp. In International conference on machine learning, 2790–2799 (PMLR, 2019).

81 Lu, X.-J., Bussemaker, H. J. & Olson, W. K. Dssr: an integrated software tool for dissecting the spatial structure of rna. Nucleic acids research 43, e142–e142 (2015).

82 Jumper, J. et al. Highly accurate protein structure prediction with AlphaFold. Nature 596, 583–589 (2021).

83 Outeiral, C. & Deane, C. M. Codon language embeddings provide strong signals for use in protein engineering. Nature Machine Intelligence 6, 170–179 (2024).

84 Nguyen, E. et al. Sequence modeling and design from molecular to genome scale with evo. Science 386, eado9336 (2024).

85. Tálas, A., et al. Bear reveals that increased fidelity variants can successfully reduce the mismatch tolerance of adenine but not cytosine base editors. Nature communications 12, 6353 (2021).

86 Danaee, P. et al. bprna: large-scale automated annotation and analysis of rna secondary structure. Nucleic acids research 46, 5381–5394 (2018).

87 Sun, S., Wang, W., Peng, Z. & Yang, J. Rna inter-nucleotide 3d closeness prediction by deep residual neural networks. Bioinformatics 37, 1093–1098 (2021).

88 Leontis, N. B. & Zirbel, C. L. Nonredundant 3d structure datasets for rna knowledge extraction and benchmarking. In RNA 3D Structure Analysis and Prediction, 281–298 (Springer, 2012).

89 Gong, J., Xu, K., Ma, Z., Lu, Z. J. & Zhang, Q. C. A deep learning method for recovering missing signals in transcriptome-wide rna structure profiles from probing experiments. Nature Machine Intelligence 3, 995–1006 (2021).

90 Jaganathan, K. et al. Predicting splicing from primary sequence with deep learning. Cell 176, 535–548.e24 (2019).

91 Bogard, N., Linder, J., Rosenberg, A. B. & Seelig, G. A deep neural network for predicting and engineering alternative polyadenylation. Cell 178, 91–106 (2019).

92 Amin, N., McGrath, A. & Chen, Y.-P. P. Evaluation of deep learning in non-coding RNA classification. Nature Machine Intelligence 1, 246–256 (2019).

93 Fiannaca, A., La Rosa, M., La Paglia, L., Rizzo, R. & Urso, A. nrc: non-coding rna classifier based on structural features. BioData mining 10, 1–18 (2017).

94 Sample, P. J. et al. Human 5’ UTR design and variant effect prediction from a massively parallel translation assay. Nat. Biotechnol. 37, 803–809 (2019).

95 Wayment-Steele, H. K. et al. Deep learning models for predicting rna degradation via dual crowdsourcing. Nature Machine Intelligence 4, 1174–1184 (2022).

96 Angenent-Mari, N. M., Garruss, A. S., Soenksen, L. R., Church, G. & Collins, J. J. A deep learning approach to programmable rna switches. Nature Communications 11, 5057 (2020). URL https://www.osti.gov/biblio/1816589.

97 Angenent-Mari, N. M., Garruss, A. S., Soenksen, L. R., Church, G. & Collins, J. J. A deep learning approach to programmable rna switches. Nature Communications 11, 5057 (2020).

98 Chuai, G. et al. Deepcrispr: optimized crispr guide rna design by deep learning. Genome biology 19, 1–18 (2018).

99 Miller, T. M., Cudkowicz, M., Shaw, P. J. et al. Phase 1–2 trial of antisense oligonucleotide tofersen for sod1 als. New England Journal of Medicine 383, 109–119 (2020).

100 Ross, S. J., Revenko, A. S., Hanson, L. L. et al. Targeting kras-dependent tumors with azd4785, a high-affinity therapeutic antisense oligonucleotide inhibitor of kras. Science Translational Medicine 9, eaal5253 (2017).

101. Beckerman, P., Bi-Karchin, J., Abed, A., et al. Antisense oligonucleotide treatment ameliorates ifn-–induced proteinuria in apol1 transgenic mice. JCI Insight 2, e92750 (2017).

102 Meng, L., Ward, A. J., Chun, S. et al. Towards a therapy for angelman syndrome by targeting a long non-coding rna. Nature 518, 409–412 (2015).

103 Mathews, D. H. How to benchmark rna secondary structure prediction accuracy. Methods 162, 60–67 (2019).

104 Reuter, J. S. & Mathews, D. H. RNAstructure: software for RNA secondary structure prediction and analysis. BMC Bioinformatics 11, 129 (2010).

105 Rives, A. et al. Biological structure and function emerge from scaling unsupervised learning to 250 million protein sequences. Proceedings of the National Academy of Sciences 118, e2016239118 (2021).

106 Zhang, Y. et al. Multiple sequence alignment-based RNA language model and its application to structural inference. Nucleic Acids Research 52, e3–e3 (2023). URL 10.1093/nar/gkad1031. https://academic.oup.com/nar/article-pdf/52/1/e3/55443207/gkad1031.pdf.

107 Boyd, N. et al. Atom-1: A foundation model for rna structure and function built on chemical mapping data. bioRxiv 2023–12 (2023).

